# Multifaceted Actions of Neurosteroids

**DOI:** 10.1101/2025.01.22.634297

**Authors:** Ajeet Kumar, Mingxing Qian, Yuanjian Xu, Ann Benz, Douglas F. Covey, Charles F. Zorumski, Steven Mennerick

## Abstract

**Background and purpose:** Neurosteroids modulate neuronal function and are promising therapeutic agents for neuropsychiatric disorders. Neurosteroid analogues are approved for treating postpartum depression and are of interest in other disorders. GABA-A receptors are well characterized targets of natural neurosteroids, but other biological pathways are likely relevant to therapeutic mechanisms and/or to off-target effects. We performed hypothesis-generating *in silico* analyses and broad *in vitro* biological screens to assess the range of actions of neurosteroids analogues of varying structural attributes.

**Key Results:** We employed *in silico* molecular similarity analysis and network pharmacology to elucidate likely targets. This analysis confirmed likely targets beyond GABA-A receptors. We then functionally screened 19 distinct neurosteroid structures across 78 targets representing interconnected signaling pathways, complemented with a limited screen of kinase activation. Results revealed unanticipated modulation of targets by neurosteroids with some structural selectivity. Many compounds-initiated androgen receptor translocation with little or no enantioselectivity. Modulation of multiple G-protein receptors was also unexpected.

**Conclusions and implications:** Neurosteroids are ascendant treatments in neuropsychiatry, but their full spectrum of actions remains unclear. This virtual and biological screening discovery approach opens new vistas for exploring mechanism of neurosteroids analogues. The multifaceted approach provides an unbiased, holistic exploration of the potential effects of neurosteroids across various molecular targets and provides a platform for future validation studies to aid drug discovery.

## INTRODUCTION

Neurosteroids (NS), synthesized endogenously within the central nervous system (CNS), modulate neuronal excitability and neurotransmission (Mellon and Griffin, 2002; Maguire and Mennerick, 2023). NS are produced by neurons and glial cells through enzymatic reactions in the cholesterol metabolic pathway, facilitated by enzymes for side-chain cleavage, aromatases, hydrolases, and reductases (Mellon et al., 2001; Stoffel-Wagner, 2001; Wang et al., 2024). They are categorized into three main groups based on their chemical structures and precursor molecules (MacKenzie and Maguire, 2013). Firstly, pregnane compounds, mainly derived from progesterone (Prog), include NS such as allopregnanolone (AlloP) and pregnanolone. Synthetic steroids such as ganaxolone, alfaxalone, and zuranolone are analogues of natural pregnane steroids. Secondly, androstane analogues, originating from testosterone, comprise steroids like androstanediol, and etiocholanone. Sulfated NS, exemplified by dehydroepiandrosterone sulfate and pregnenolone sulfate, constitute another significant category in this structural classification scheme. The mode of action of NS involves interaction with specific neurotransmitter receptors (Akk et al., 2007) but also other targets. Prominent among those that interact with transmitter receptors are AlloP and similar pregnane neurosteroids. These compounds show therapeutic potential in various neuropsychiatric disorders (Zorumski and Mennerick, 2013). Brexanolone (AlloP) and zuranolone (a 5β-reduced pregnane analogue) have recently gained FDA approval as rapid-acting antidepressants specifically for treating postpartum depression (Maguire and Mennerick, 2023; Patterson et al., 2023). Ganaxolone, another pregnane analogue, is approved for the treatment of seizures associated with cyclin-dependent kinase-like 5 (CDKL5) deficiency disorder (CDD) (Keam and Al-Salama, 2023; Keam et al., 2023).

Endogenous NS and their analogues exert key effects by interacting with gamma-aminobutyric acid type A (GABA-A) receptors, the principal fast inhibitory neurotransmitter receptors in the CNS (Lambert et al., 2001; Belelli and Lambert, 2005; Farrant and Nusser, 2005; Belelli et al., 2021; Legesse et al., 2023). Pregnane NS function as positive allosteric modulators (PAMs). By binding to GABA-A receptors, NS PAMs enhance the inhibitory actions of GABA, resulting in increased influx of chloride ions into neurons (Belelli and Lambert, 2005; Tuem and Atey, 2017; Zorumski et al., 2019) to reduce neuronal excitability. This heightened inhibition may contribute significantly to a role for NS in regulating mood, anxiety, stress responses, and other CNS processes (Mellon and Griffin, 2002; MacKenzie and Maguire, 2013; Zorumski et al., 2013). At high concentrations, pregnane steroids also appear to inhibit GABA-A receptor function (Hauser et al., 1996; Foll et al., 1997; Zhu and Vicini, 1997) although the physiological relevance is unclear. Despite the well characterized effects of NS on inhibition, not all GABA-A PAMs (e.g., propofol, barbiturates) appear to have the same neuropsychiatric benefit. Further, NS has side effects clinically that limit their use (Handan et al., 2018; Handan and Kanes, 2019; Kleinman and Schatzberg, 2021). These observations suggest that targets other than GABA-A receptors should be considered in the quest to improve these medications.

Several preclinical and clinical studies suggest that NS interacts with various other proteins and receptors (Vallée et al., 2014; Irina et al., 2019; Mosher et al., 2019; Balan et al., 2023). In addition to ion channels and G protein-coupled receptors (GPCRs), some studies suggest that NS indirectly modulates kinase activity (Adams et al., 2000; András and Linn, 2000; Modgil et al., 2017; Parakala et al., 2019; Littlejohn and Boychuk, 2021). Thus, research is needed to uncover unorthodox protein targets for NS, potentially advancing novel therapeutic modalities. Here we assessed the molecular similarity of NS with small molecules from open-source databases to assess targets by drug-target interaction. This approach has been used previously (Janssen et al., 2019; Li et al., 2020). Furthermore, we used network pharmacology with drug targets to determine the classes of targets that are strongly associated with NS like molecules. This method is also previously validated (Duran-Frigola et al., 2020; Tanoli et al., 2020; Nuzzo et al., 2021; Patil et al., 2024).

Based on molecular similarity results and NS target network analysis, we hypothesized that NS could modulate several targets other than GABA-A receptors. In this study, we investigated nineteen NS and analogues of NS from pregnane, androstane, and sulfate-like categories (e.g. AlloP, Prog, ganaxolone, alfaxalone, MQ34, B260, CDCN24, etiocholanolone (Etio), ECN, 3α5βPC and androsterone (Andro)), including four synthetic enantiomers (*ent*-AlloP, *ent*-Prog, *ent*-Etio, and MQ35). Subsequently, we evaluated biological activity at targets identified through network pharmacology analysis and through the availability of commercial screening assays. Targets encompassed neurotransmitter GPCRs, ion channels, and enzymes. We confirmed the biological activity of NS toward acetylcholine receptors (Paradiso et al., 2001) and identified previously unknown interactions with HTR3A serotonergic receptors (HTR3A), voltage-gated channels (Na_v_1.5) and beta-adrenergic receptors related to cardiac functions. Although some of these effects are restricted to high NS concentrations and caution is warranted by some mismatches in screening results with known NS actions in the CNS, our hypothesis-generating results underscore diverse effects of NS at pharmacological concentrations that are relevant to their potential clinical use.

## METHODS

### Availability of material, data, and code

This study did not generate any new large data sets. All screening related data sets are available in supplementary files, and small molecule library data sets are available open source for academic research use.

Raw data reported in this paper will be shared by the corresponding author upon reasonable request. Python analysis scripts were developed with the assistance of open-source resources and OpenAI tools to efficiently organize, process, and analyze data. Any additional information required to reanalyze the data reported in this paper is available from the corresponding author upon request.

### Reagent sources

| REAGENTS | SOURCE | IDENTIFIER/<br>REFERENCE |
| --- | --- | --- |
| **Commercial assays** |  |  |
| DiscoverX<br>HitHunter cAMP<br>XS+ assay | Eurofins Discovery | https://www.eurofinsdiscovery.com/about-us |
| PathHunter®NHR<br>Protein Interaction<br>Assays | Eurofins Discovery | https://www.eurofinsdiscovery.com/about-us |
| PathHunter®<br>Nuclear<br>Translocation<br>assays | Eurofins Discovery | https://www.eurofinsdiscovery.com/about-us |
| PathHunter®<br>Calcium No<br>Wash<sup>PLUS</sup> assay | Eurofins Discovery | https://www.eurofinsdiscovery.com/about-us |
| FLIPR®<br>Potassium Assay | Eurofins Discovery | https://www.eurofinsdiscovery.com/about-us |
| FLIPR®<br>Membrane<br>Potential Assay | Eurofins Discovery | https://www.eurofinsdiscovery.com/about-us |
| DiscoverX<br>Calcium<br>NWPLUS Assay | Eurofins Discovery | https://www.eurofinsdiscovery.com/about-us |
| Neurotransmitter<br>Transporter<br>Uptake Assay | Eurofins Discovery | https://www.eurofinsdiscovery.com/about-us |
| KINOMEScan® Assay | Eurofins Discovery | https://www.eurofinsdiscovery.com/about-us |
| Adapta® Universal Kinase Assay | Thermo Fisher Sci | www.thermofisher.com/selectscreen |
| **Chemicals** |  |  |
| Allopregnanolone | Sigma Aldrich | P8887 |
| *ent*-Allopregnanolone | Covey Lab | (Hu et al., 1997) |
| Progesterone | Sigma Aldrich | P8783 |
| *ent*-Progesterone | Covey Lab | (Auchus et al., 2003) |
| Androsterone | Sigma Aldrich | 31579 |
| *ent*-Androsterone | Covey Lab | (Katona et al., 2007) |
| Etiocholanolone | Sigma Aldrich | E5126 |
| *ent*-Etiocholanolone | Covey Lab | (Katona et al., 2007) |
| Alfaxalone | Covey Lab | (Tescic et al., 2020) |
| Ganaxalone | Covey Lab | (Hogenkamp et al., 1997) |
| 3α5βPC | Covey Lab | (Mennerick et al., 2001) |
| CDNC24 | Covey Lab | (Tescic et al., 2020) |
| ECN | Covey Lab | (Todorovic et al., 1998) |
| B260 | Covey Lab | (Han et al., 1996) |
| B372 | Covey Lab | (Han et al., 1996) |
| MQ34 | Covey Lab | (Qian et al., 2014) |
| MQ35 | Covey Lab | (Qian et al., 2014) |
| MQ221 | Covey Lab | (Ziolkowski et al., 2020) |
| YX23 | Covey Lab |  |
| **Software and algorithms** |  |  |
| Python |  | 3.12 |
| Jupyter |  | https://jupyter.org/ |
| RDKit |  | https://www.rdkit.org/ |
| ChemDraw |  | 22.2.0 |
| GraphPad Prism |  | 10.4.0 |
| **Database** |  |  |
| Drug Repurposing Hub | Broad Institute | https://www.broadinstitute.org/drug-repurposing-hub |
| DrugBank | DrugBank 5.1.12 | https://go.drugbank.com/releases/latest |
| ChEMBL library | ChEMBL_webresource | https://www.ebi.ac.uk/chembl/ |
| STRING Library | version 12.0 | https://string-db.org/ |

### Neurosteroids synthesis

AlloP, Prog, Etio and Andro were purchased from Sigma Aldrich and other NS and analogues used in screening were synthetized in the laboratory of Douglas F. Covey. Synthetic details are provided in references given in the Identifier/Reference column of the Key Resource Table. Reviews of synthetic methods have also been published (Jastrzębska and Covey, 2022; Covey et al., 2023).

### Data collection and molecular similarity search

In this survey, we collected small molecule data from three openly accessible databases. Specifically, we obtained 6798 drugs molecules including investigational drugs molecule from the Drug Repurposing Hub, along with their respective target names available at http://www.broadinstitute.org/repurposing. Additionally, we downloaded 11926 small molecules and their associated target names from the DrugBank library (https://go.drugbank.com), as well as 3492 approved drugs along with their targets from the ChEMBL library, utilizing web resources through Python scripts ([CSL STYLE ERROR: reference with no printed form.]). The canonical SMILES (Simplified Molecular Input Entry System) representations of these molecules were extracted using a Python program integrated with the open-source cheminformatics tool RDKit.

To identify neurosteroid-like molecules, we used t-distributed stochastic neighbor embedding (t-SNE) maps for a molecular similarity search in the chemical space of approved drugs and investigational small molecules. This method was previously used in several published articles (Janssen et al., 2019; Duran-Frigola et al., 2020; Li et al., 2020). AlloP, an FDA-approved NS, was used as query molecule for a similarity search in chemical space. The subsequent step generated Morgan fingerprints for each molecule using RDKit’s “GetMorganFingerprintAsBitVect” function and the parameters radius=2 and nBits=4096. The radius parameter dictates the breadth of the fingerprint’s neighborhood, influencing the level of detail captured in the molecular representation. Meanwhile, the nBits parameter determines the length of the fingerprint bit vector, affecting the granularity of the molecular features encoded. These parameters strike a balance between capturing sufficient structural information and managing computational complexity. Following fingerprint generation, t-SNE was employed to project the high-dimensional molecular space into a more manageable two-dimensional representation.

The parameters, components=2 and random state=42, were specified to indicate the desired number of dimensions in the reduced space and ensure reproducibility across runs, respectively. By preserving local structure and revealing underlying patterns, t-SNE enabled intuitive exploration and interpretation of the molecular dataset. Subsequently, k-nearest neighbors (k-NN) was applied with parameter k=50 to determine the number of neighbors considered in the search. By computing distances between the query molecule and its nearest neighbors in the fingerprint space, this step facilitated the identification of molecules sharing structural similarities, thus offering valuable insights into potential drug candidates related to the query molecule. In parallel, K-means clustering was employed to partition the dataset into distinct groups based on the t-SNE-transformed features. To provide a visual reference for molecules within a certain distance from the query, a circle with a radius of 5 units was plotted around the query molecule in the t-SNE visualization. Additionally, extracted information for molecules within the circle surrounding the query molecule was obtained. This information typically included details such as Molecule Name, Mechanism of Action (MOA) and Target names. This information provided characteristics of molecules closely related to AlloP, aiding in further analysis and interpretation.

### Drug-target network pharmacology

We conducted a similarity search to identify neurosteroid-like molecules and their targets with associated genes, followed by manual verification of target names using additional sources. In cases where target names were similar for multiple molecules, we ensured selection of each target only once for network pharmacology analysis. All potential target genes shared among the molecules were integrated into the STRING version 12.0 database (https://string-db.org) to construct a protein-protein interaction (PPI) network, with Homo sapiens selected as the organism. We opted for a medium confidence level, setting an interaction score threshold of >0.4 for the PPI network. This network consists of nodes representing target proteins and edges representing protein-protein interactions. The degree of each node indicates the number of direct connections it possesses, with higher degrees indicating greater importance.

### GO and KEGG enrichment analysis

Gene Ontology (GO) and KEGG (Kyoto Encyclopedia of Genes and Genomes) pathways analyses are essential components of drug-target identification and development, playing integral roles in understanding disease mechanisms, identifying therapeutic targets, and advancing drug discovery efforts. The STRING database also offers functionalities for functional enrichment analysis, including KEGG and GO analysis. In this analysis we used similar data sets as used in network pharmacology, KEGG pathway and functional enrichment of GO including biological process, molecular function and cellular component results downloaded and further analyzed with python script for visualization purposes. Top 50 enriched KEGG pathways, GO-term based on the observed-to-background gene count ratio, with each bar color-coded by the false discovery rate (FDR) value.

### Target selection

Following enrichment analysis, we meticulously scrutinized each pathway and GO-term, weighing numerous factors to choose targets for experimental validation. Our primary criterion was to prioritize targets aligned with our research focus, especially those yet unexplored for our NS molecules. We identified targets significantly enriched in pertinent GO-terms or KEGG pathways target genes, indicating their potential pivotal roles in the biological system with therapeutic implications. Moreover, we analyzed the interactions among these targets, recognizing highly connected ones as potential hubs or key regulators within biological networks. We evaluated the functional significance of the targets based on their roles in biological pathways, gene ontology categories, or protein-protein interaction networks. Although some targets lacked multiple connections, they were novel to our study. We conducted a literature review to gather further evidence supporting the involvement of selected targets in relevant biological processes or diseases. Additionally, we considered the feasibility of experimental validation, considering factors such as experimental techniques and required resources.

### *In vitro* screening

Broad *in vitro* biological screenings described below were conducted by Eurofins DiscoverX Corporation (11180 Roselle Street, Suite D, San Diego, CA 92121). We outsourced all *in vitro* screening for ion channels, GPCRs, nuclear receptors, transporters, non-kinase enzymes, and some kinases using SAFETYscan E/IC50 ELECT service from Eurofins. We also screened single doses (10 µM) of AlloP, Prog, and their enantiomers for kinase inhibition activity across 481 kinases using SelectScreen Kinase Profiling Services from Thermo Fisher (www.thermofisher.com/selectscreen).

### Ion channel assays

For ion channel activity assays, including ionotropic receptors, cell lines were seeded into Poly-D-lysine-coated, black-walled, clear-bottom 384-well microplates, followed by incubation at 37°C. Dye loading used a 1X Dye loading buffer with 2.5 mM Probenecid, incubated for 30-60 minutes at 37°C. For agonist determination, cells were incubated with the test sample, while antagonist determination involved pre-incubation with the sample in EC_80_ agonist. Signal detection was performed on a FLIPR Tetra, and data analysis used the CBIS suite to calculate percentage activity or inhibition based on relative luminescence units (RLU).

### GPCR cAMP assay

cAMP modulation was evaluated using the DiscoverX HitHunter cAMP XS+ assay. In the Gs agonist format, cells were treated with the sample, the media was replaced with HBSS/Hepes and cAMP XS+ Ab reagent, followed by incubation. For the Gi agonist format, cells were exposed to the sample in the presence of EC_80_ forskolin, with subsequent steps similar to the Gs agonist protocol. In the antagonist format, cells were first pre-incubated with the sample, then challenged with an agonist, followed by media replacement and further incubation. Signal detection was performed using cAMP XS+ reagents, with readings obtained via a PerkinElmer Envision™ instrument. Data analysis was completed using the CBIS data analysis suite, applying specific formulas to determine percentage activity or inhibition depending on the assay mode.

### Calcium flux assay

Cells were seeded into Poly-D-lysine coated 384-well microplates and incubated before testing. Assays were conducted in a 1X dye loading buffer containing dye, Additive A, and Probenecid. Cells were loaded with dye, incubated, and then tested for agonist or antagonist activity using FLIPR Tetra. Agonist activity was measured by adding a compound and monitoring calcium mobilization, while antagonist activity was measured after pre-incubation with a sample and subsequent agonist challenge. Data analysis involved calculating the area under the curve and determining percentage activity or inhibition using specific formulas.

### Nuclear translocation assay for nuclear receptor

PathHunter nuclear hormone receptor (NHR) cell lines were seeded into white-walled 384-well microplates containing charcoal-dextran filtered serum. To determine agonist activity, cells were treated with a 5X sample and incubated for 3-16 hours. For antagonist determination, cells were pre-incubated with an antagonist, then challenged with an EC_80_ agonist and incubated. Signal detection was achieved by adding the PathHunter Detection reagent cocktail and measuring the chemiluminescent signal using a PerkinElmer Envision™ instrument. Data analysis involved calculating percentage activity for agonists and percentage inhibition or inverse agonist activity for antagonists, with results capped at 0% or 100% as necessary

### KINOMEscan binding assays

A kinase-tagged T7 phage was grown in E. coli, followed by infection and incubation until lysis; lysates were centrifuged and filtered, and kinases were produced in HEK-293 cells and tagged for qPCR detection. Streptavidin-coated beads were prepared with biotinylated ligands and blocked to create affinity resins, which were combined with kinases, liganded beads, and test compounds in binding reactions. After incubation, kinase concentrations were measured by qPCR. Percent response and binding constants (Kds) were calculated using a dose-response curve.

### Transporter assays

Transporter cell lines were seeded into Poly-D-lysine coated 384-well microplates and incubated at 37°C prior to testing. After cell plating and incubation, the media was removed, and 25 µL of 1X compound in HBSS/0.1% BSA was added, followed by a 30-minute incubation at 37°C. Assays were performed in 1X Dye Loading Buffer, with 25 µL of 1X dye added to the wells, and cells were incubated for 30-60 minutes at 37°C. Fluorescence signal detection was then conducted using a PerkinElmer Envision™ instrument, and compound activity was analyzed using the CBIS data analysis suite, where percentage inhibition for blocker mode assays was calculated, and percent response for primary screens was capped at 0% or 100% based on the results.

### Enzymatic assays

After preparing enzymes, including AChE, COX1, COX2, MAOA, PDE3A, and PDE4D2, enzymatic activity assays were conducted to measure substrate consumption or product formation over time using various detection methods. For AChE, the enzyme and test compounds were preincubated before substrate addition, with the signal detected by absorbance at 405 nm. For COX1 and COX2, the enzymes were equilibrated with compounds, and the reactions were measured on a fluorimeter. For MAOA, reactions were initiated with kynuramine and terminated with NaOH, with signal detection via spectrofluorimetry(Matsumoto et al., 1985). For PDE3A and PDE4D2, the reactions involved cAMP substrate and were terminated by 3-isobutyl-1-methylxanthine (IBMX), with detection using a cAMP detection kit. Signal detection was performed using a PerkinElmer Envision™ instrument, and data were analyzed using the CBIS data suite to calculate percentage inhibition and percent response in primary screens.

### Analyses

We used GraphPad Prism 10 for dose response curve fit analysis to generate EC_50_/IC_50_ of all NS analogues. We used Python based Matplotlib for visualization of data, such as heatmap, t-SNE plot and gene enrichment analysis.

## RESULTS

### Molecular similarity of neurosteroids with investigational & approved drugs

To explore novel therapeutic targets of NS, we started with a cheminformatics approach. We compared chemical space similarity of NS with other bioactive small molecules from open-source, small-molecule databases, using AlloP as the query molecule because it is a clinically approved NS prototype. In the resulting t-distributed stochastic neighbor embedding (t-SNE) map with Drug Repurposing Hub molecules (Figure 1A), each point is a molecule, colored according to its assigned cluster. The query molecule (AlloP) is highlighted as a red cross in the plot, facilitating visual inspection of its neighborhood in the reduced-dimensional space. Unsurprisingly, many near-neighbors had a familiar steroid-like backbone (Figure 1B).

**Figure 1.**
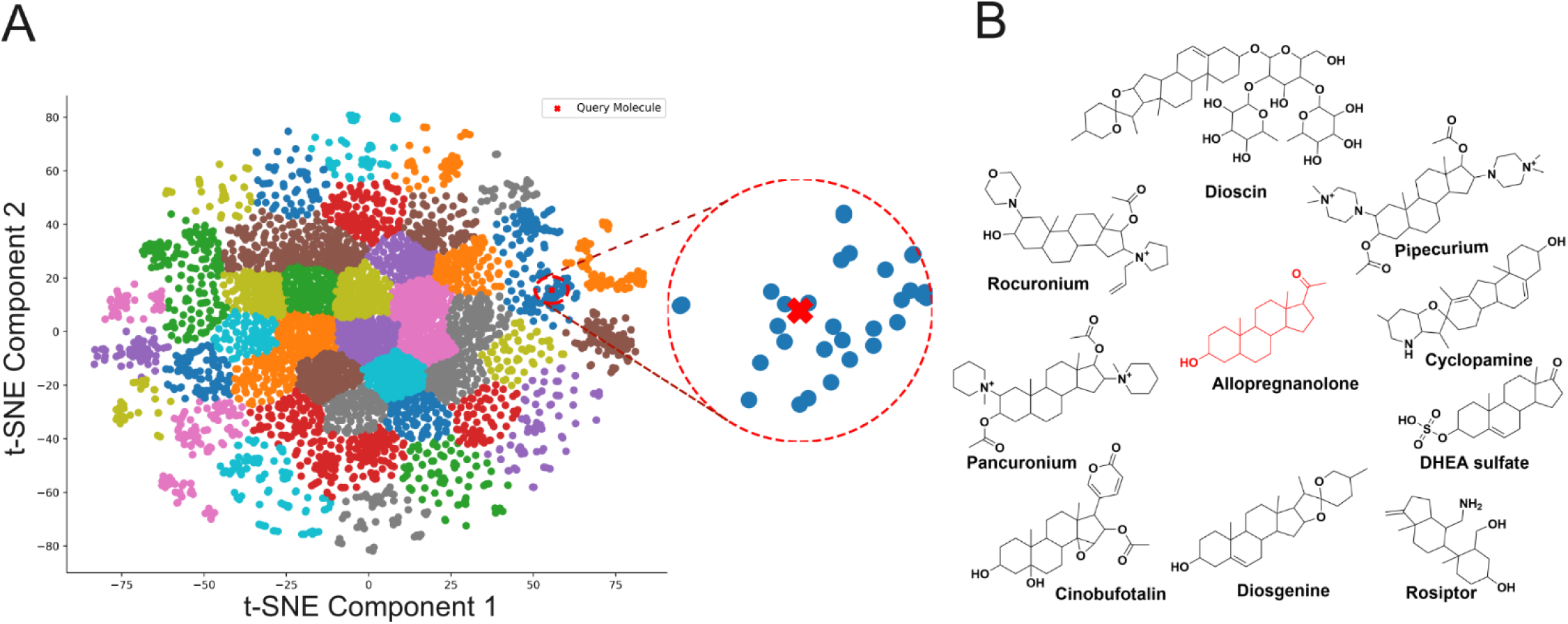
t-SNE based chemical similarity analysis of AlloP with the Drug Repurposing Hub dataset and selected similar molecules. **(A)** t-SNE visualization of Drug Repurposing Hub dataset with query molecule (AlloP) in 2D chemical space. The analysis was performed using 4096-bit Morgan fingerprinting with default t-SNE setting for embedding. Molecular similarity was determined by k-nearest neighbors’ analysis and chemical space distance from AlloP. A circle with a radius of 5 units was plotted around the query molecule in the t-SNE visualization. This analysis helped identify potential drug candidates with comparable chemical properties to AlloP, suggesting possible shared biological activities. **(B)** Selected similar molecules with a steroid-like scaffold are depicted alongside AlloP, to demonstrate shared chemical features.

Upon performing the t-SNE based chemical similarity search from Drug Repurposing Hub molecules (6798 drugs molecules), we investigated 37 neighboring molecules located within a 5-unit radius (red dotted circle) of chemical space around AlloP (Figure 1A and 1B). All neighboring molecules are listed in Table 1 along with varied modes of action of the drugs. These molecules showed a wide range of pharmacological activities, including androgen receptor modulation, anticancer activity, inhibition of the hedgehog pathway, and modulation of neurotransmitter receptors. Neurotransmitter receptor targets included GABA-A, glutamate receptors and both muscarinic and nicotinic acetylcholine receptors. Other targets included an ionotropic serotonin receptor (HTR3A), chemokine receptor (CXCR3), smoothened GPCR, and kinase associated proteins (Src, FAK, Erk, PI3K, mTOR, GSK3, CDKL5, SHIP1). The queried molecules have a steroidal backbone and generally, are known for their significant activity towards nuclear receptors including estrogen receptors (ESR1, ESR2), androgen receptor (AR) and glucocorticoid receptor (GR). The molecules are also involved in the activity of cytochrome P450 enzymes (CYP19A1, CYP17A1), and some show NS-like modulation of GABA-A receptors (GABRA1-6, GABRB1-3, GABRD-RE) and glutamate receptors (GRIN1, GRIN2A-B, GRIN3A-B).

**Table 1.**
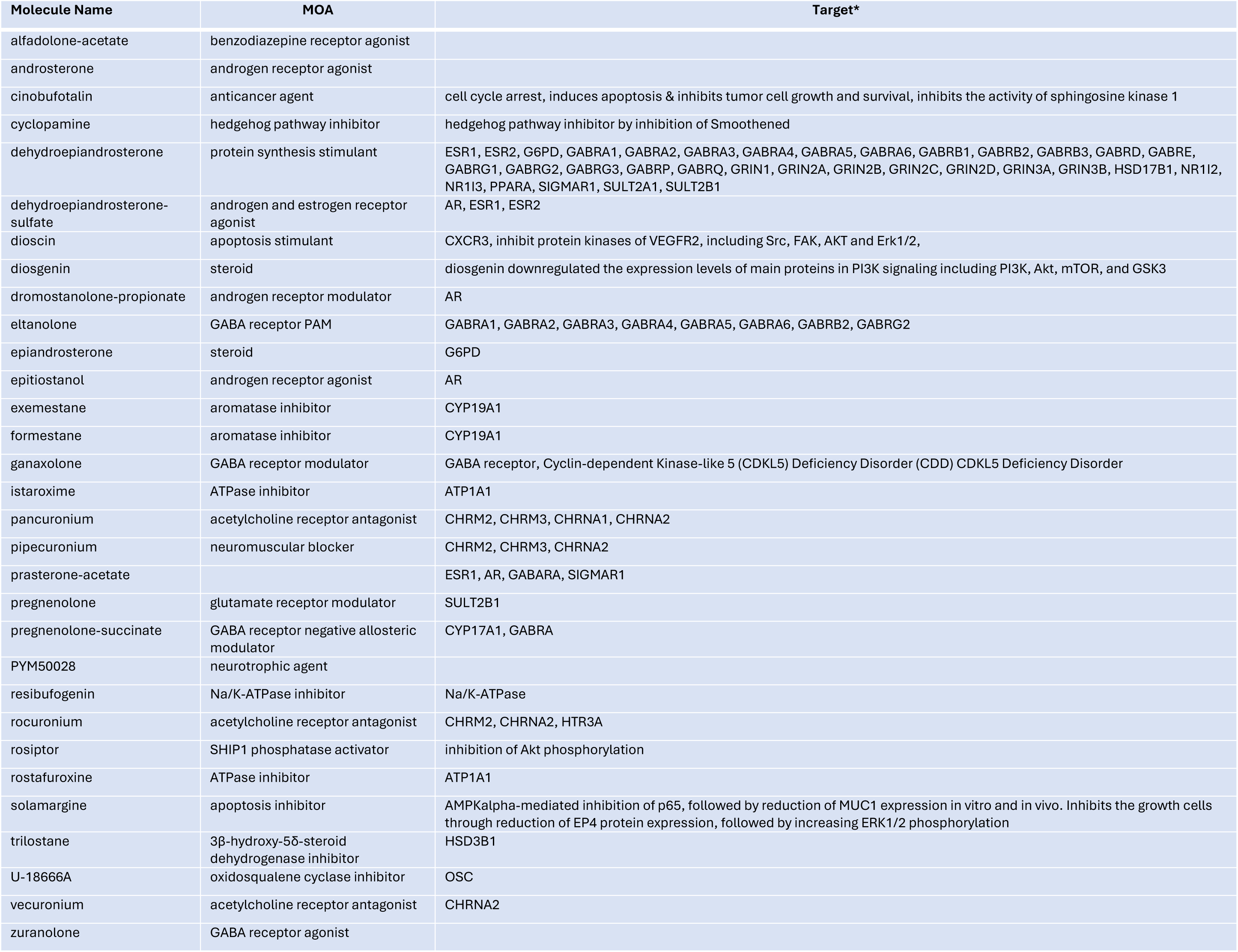
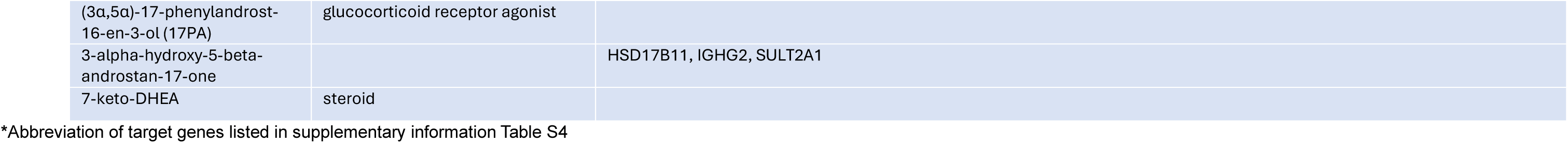
Neighboring molecules and reported mechanisms of action.

In addition to the Drug Repurposing Hub database, we also utilized the DrugBank and ChEMBL datasets for chemical similarity searches. Analysis of the DrugBank dataset through t-SNE embedding (Figure S1) revealed a similar list of molecules (Table S1) to those in the DrugRepurposingHub data set. From the ChEMBL dataset, we used approved drug molecules, which retrieved only four molecules including the query molecule allopregnanolone (Figure S2 & Table S2). This analysis demonstrates that AlloP has few similar approved drugs. Notably, the neighboring molecules of AlloP from the Drug Repurposing Hub dataset encompass targets across all classes found in DrugBank and ChEMBL. Hence, we restricted our further analysis to the targets within the Drug Repurposing Hub dataset.

### Network pharmacology and enrichment analysis

The targets identified from the DrugRepurposingHub dataset offer insights into drug action, safety profile, and therapeutic potential, and contribute to the identification of new targets for NS. To explore the most important targets among the set, we employed protein-protein interaction (PPI) network analysis and enrichment analysis using the STRING database (Figure 2). This analysis shows connections weighted by the degree of published exploration, so thin connections may reveal important but relatively under-investigated pathways. Our network analysis elucidated lists of pivotal targets such as GABA and glutamate receptors, steroid synthesis and signaling, but it also revealed other novel targets such as acetylcholine GPCRs (CHRM2 and CHMR3), nicotinic acetylcholine receptors, serotonin receptors, and kinases (Figure 2A). These novel targets offer a future investigation to explore NS bioactivity beyond the well-studied GABA-A receptors.

**Figure 2.**
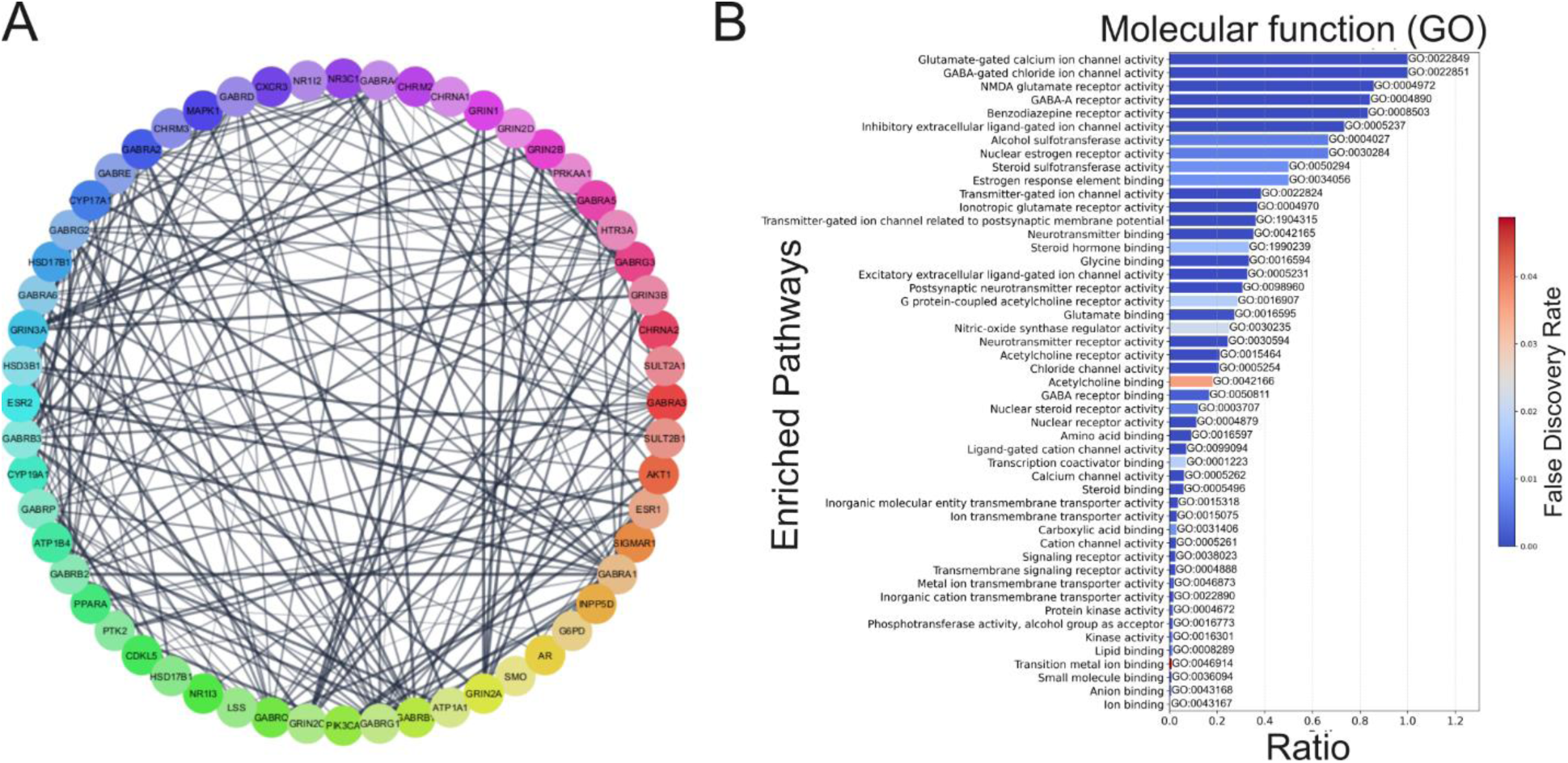
Drug target gene network and Gene Ontology (GO) enrichment analysis of molecular function. **(A)** Drug target gene network of neighbor molecules generated using the STRING database. Nodes represent individual genes, and edges denote predicted or known protein-protein interactions among these genes. The network highlights the interconnected nature of the drug targets and their potential functional associations. **(B)** Drug target gene ontology (GO) enrichment analysis of the network identified significantly overrepresented molecular functions associated with the drug target genes. These enriched functions are depicted in the associated bar chart, with each bar representing a distinct molecular function category, ranked by significance false discovery rate and ratio of number of genes associated with a particular molecular function to the total number of genes.

Enrichment analysis helps identify overrepresented biological terms or pathways among the target proteins, elucidating their functional significance. In this study, we conducted gene ontology (GO) enrichment analysis to explore the molecular functions of AlloP neighbor targets (Figure 2B). Among the top 50 significantly enriched molecular functions are GABA receptor activity, glutamate receptor activity and nuclear steroid receptor activity, which are well-known for their involvement with steroid-like molecules (Figure 2B). The analysis revealed other important molecular functions covering multiple neurotransmitter receptors and signaling systems. Many of these have been explored less than GABA-A receptors for NS activity, and we selected several for our study of the polypharmacology of NS molecules.

### Biological activity of neurosteroid analogues

Based on molecular function results, we chose targets to examine for *in vitro* activity of our natural and synthetic NS molecules (Figure 3). We evaluated NS activity at ionotropic glutamate receptor (NMDAR1A/2B), neurotransmitter receptors including GPCRs (muscarinic: CHRM1-3, serotonergic: HTR1-2, dopaminergic: DRD1-2, histaminergic: HRH1-2, adrenergic: ADRA1-2, ADRB1-2), other transmembrane receptor (adenosine: ADOA2A, arginine vasopressin: AVPR1A, cholecystokinin: CCKAR, cannabinoid: CNR1-2, opioids: OPRD1, OPRK1, OPRM1 and insulin: INSR), ligand gated ion channel (serotonergic: HTR3A and nicotine acetylcholine: nAChR(α4/β2), calcium ion channel (Ca_v_1.2), and nuclear receptors (AR and GR). We also included neurotransmitter associated transporters; serotonin (SERT), dopamine (DAT) and norepinephrine (NET) and enzymes; acetylcholinesterase (AChE) and monoamine oxidase (MAOA) that were highly involved in neurotransmitter activity functions. Despite kinase activity being situated near the bottom of our enrichment analysis (Figure 2B), we selected kinase targets for activity evaluation due to their novelty and potential functional relevance; these were tested with a subset of NS analogues at a single concentration. Some targets were also included that did not appear in the enrichment analysis but have strong relevance to side effects or were part of a commercial screening package; examples include KvLQT1, hERG, Cox1-2, EDNRA, PDE3A and PDE4D2.

**Figure 3.**
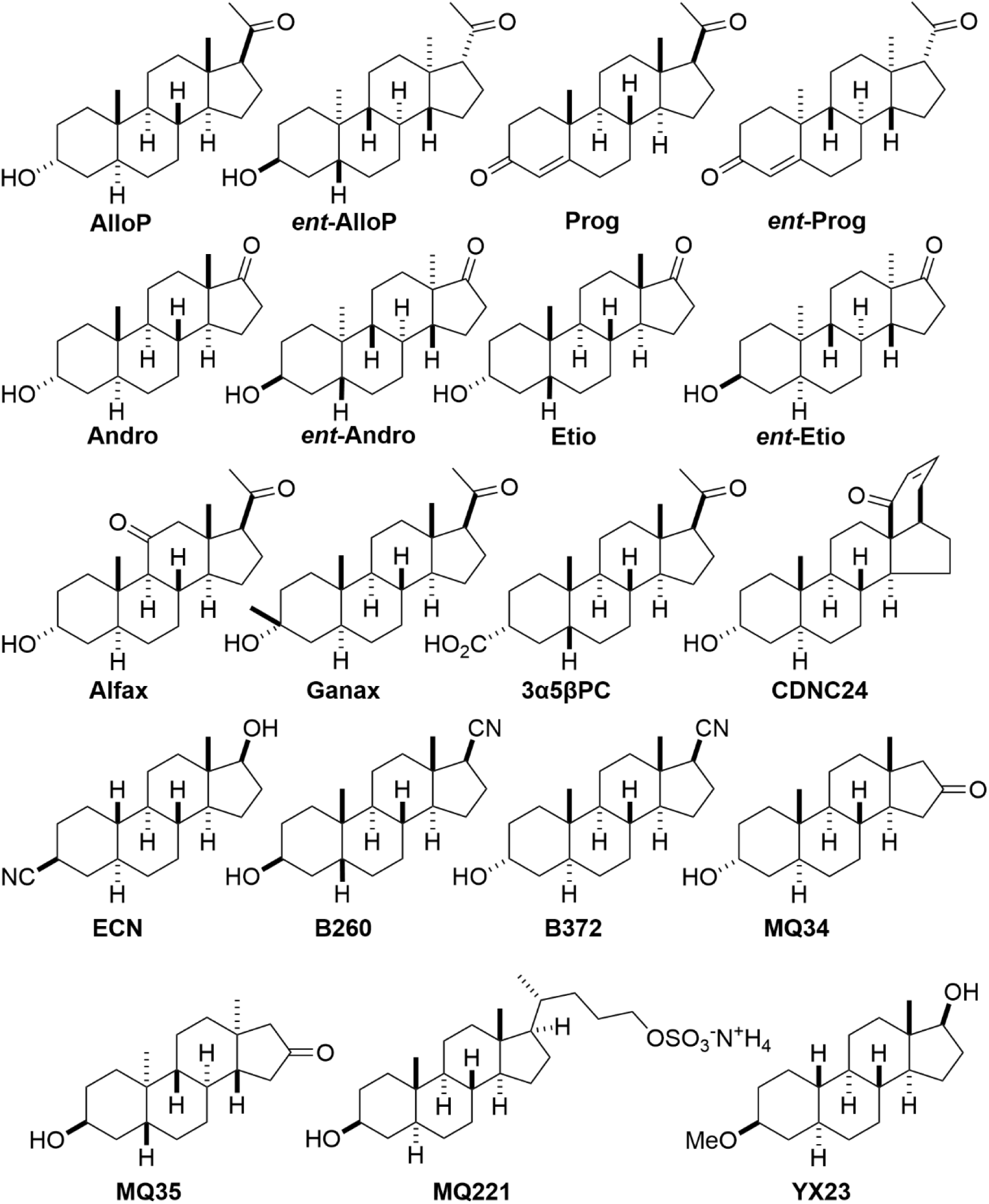
Chemical structure of neurosteroids and their analogues. The figure illustrates the chemical structures of various neurosteroids and their analogues used in the *in vitro* screening at various targets to determine IC_50_/EC_50_.

All 19 NS and analogues (Figure 3) were tested for activity in a concentration dependent manner (1.5 nM to 30 µM) to determine EC_50_/IC_50_ values at chosen targets. To present results in a compact form, we represented all tested NS activity in heatmaps (Figure 4, 5 and 6), EC_50_/IC_50_ values in Table 2 and Table S3, and we discuss the activity results in the following sections.

**Figure 4.**
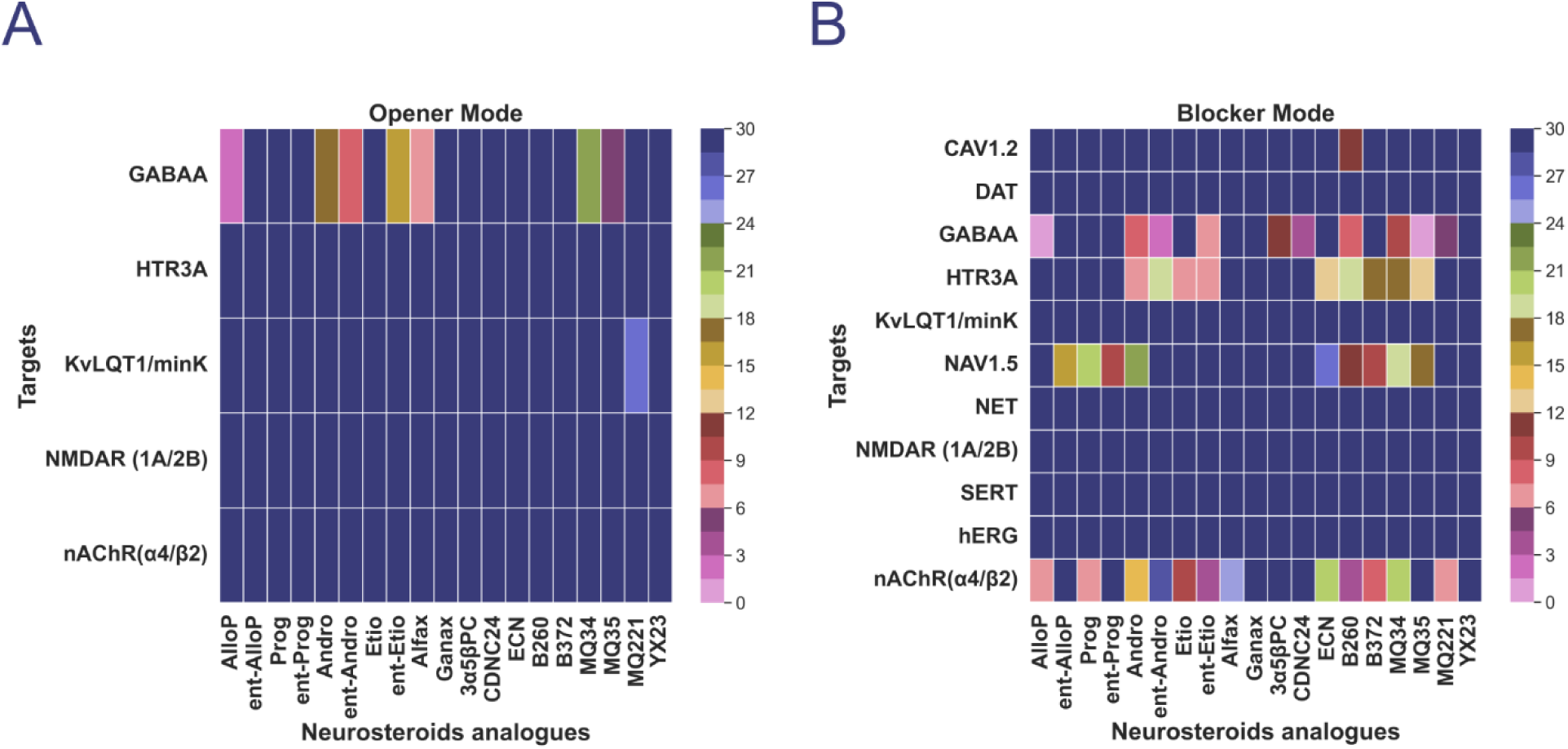
Heatmap of in-vitro activity of neurosteroids and their analogues at various ion channel and transporter targets. **(A)** The heatmap represents the agonist activity profiles of a series of neurosteroids and their analogues across multiple ion channel targets in opener modes. **(B)** The heatmap represents the blocker mode activity profiles across multiple ion channel targets and neurotransmitter transporter. Scale bar color map transitions from darker shades (representing lower activity) to lighter shades (indicating higher activity). Custom tick marks are defined at intervals of 3 µM units, ranging from 0 to 30 µM.

**Table 2.**
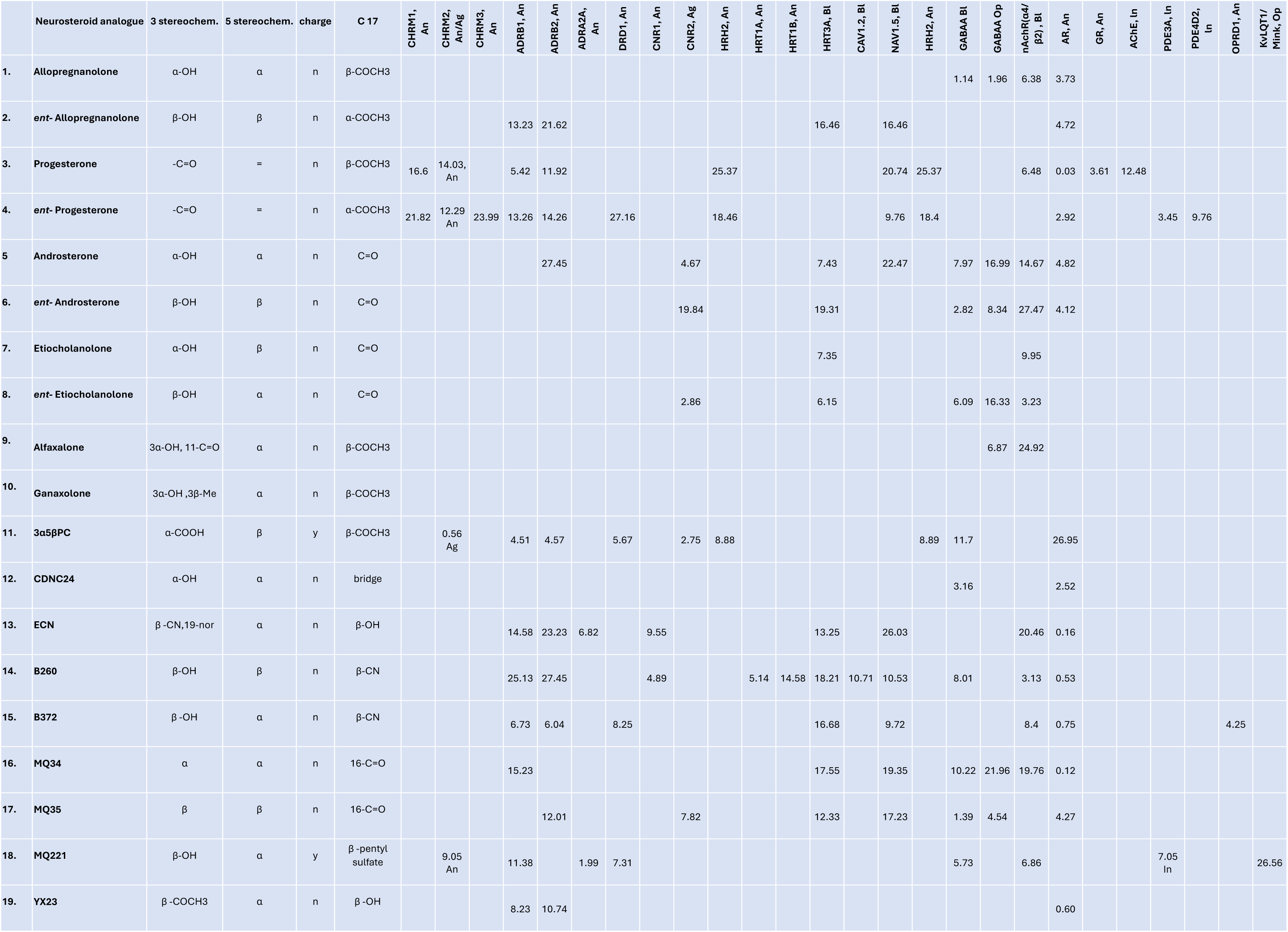
Key structural attributes of NS analogues and screening outcomes. Ag= agonist, An= antagonist. Values are EC50/IC50 values in µM.

### Cholinergic receptor activity

Acetylcholine receptors have two classes, ionotropic nicotinic receptors (nAChR) and metabotropic muscarinic receptors (Picciotto et al., 2012). We evaluated NS activity at nicotinic acetylcholine α4β2 receptors (nAChRα4β2) using the FLIPR Tetra System in opener and blocker format (Figure 4A and B). Among the nineteen NS analogues, twelve exhibited blocker activity at nicotinic α4β2 receptors in a dose dependent manner (Figure 4B and S4A), and six did not show activity (Figure 4A). AlloP exhibited an IC_50_ value of 6.38 µM, similar to Prog (IC_50_ 6.48 µM) and the sulphated NS analogue MQ221 (IC_50_ 6.86 µM), while the unnatural enantiomers of AlloP and Prog did not show any activity (Figure 4B). B260, with a carbonitrile group at carbon 17 (C17), had the highest potency (IC_50_=3.13 µM), and an unnatural synthetic enantiomer of Etio, with a ketone group at C17, exhibited similar potency (IC_50_=3.23 µm). By contrast, the 5α diastereomer of B260 (B372) and natural Etio had similar but higher IC_50_ values of 8.4 µm and 9.95 µM respectively. Other NS analogues including Andro, *ent*-Andro, alfaxalone, ECN and MQ34 exhibited low potency (14 to 27 µM).

At muscarinic acetylcholine receptors, the calcium flux assay for CHRM1/3 (Gαq coupled) and cAMP assay for CHRM2 (Gαi/o coupled) were employed in agonist and antagonist modes (Figure 5A and B). Among the molecules tested, only 3α5βPC showed potent agonist activity (EC_50_=0.56 µM) at CHRM2 in the cAMP assay in a concentration dependent manner (Figure S4C). Conversely, analogue MQ221 only showed antagonistic activity toward CHRM2 (IC_50_= 9.05 µM), while Prog and *ent*-Prog exhibited antagonism with IC50 values ranging from 12 µM to 22 µM at CHRM1 and CHRM2 (Figure S4B, and S4D). Finally, *ent*-Prog also exhibited low-potency antagonism of CHRM3 with IC50 value 23.99 µM (Figure 4B; Table 2).

**Figure 5.**
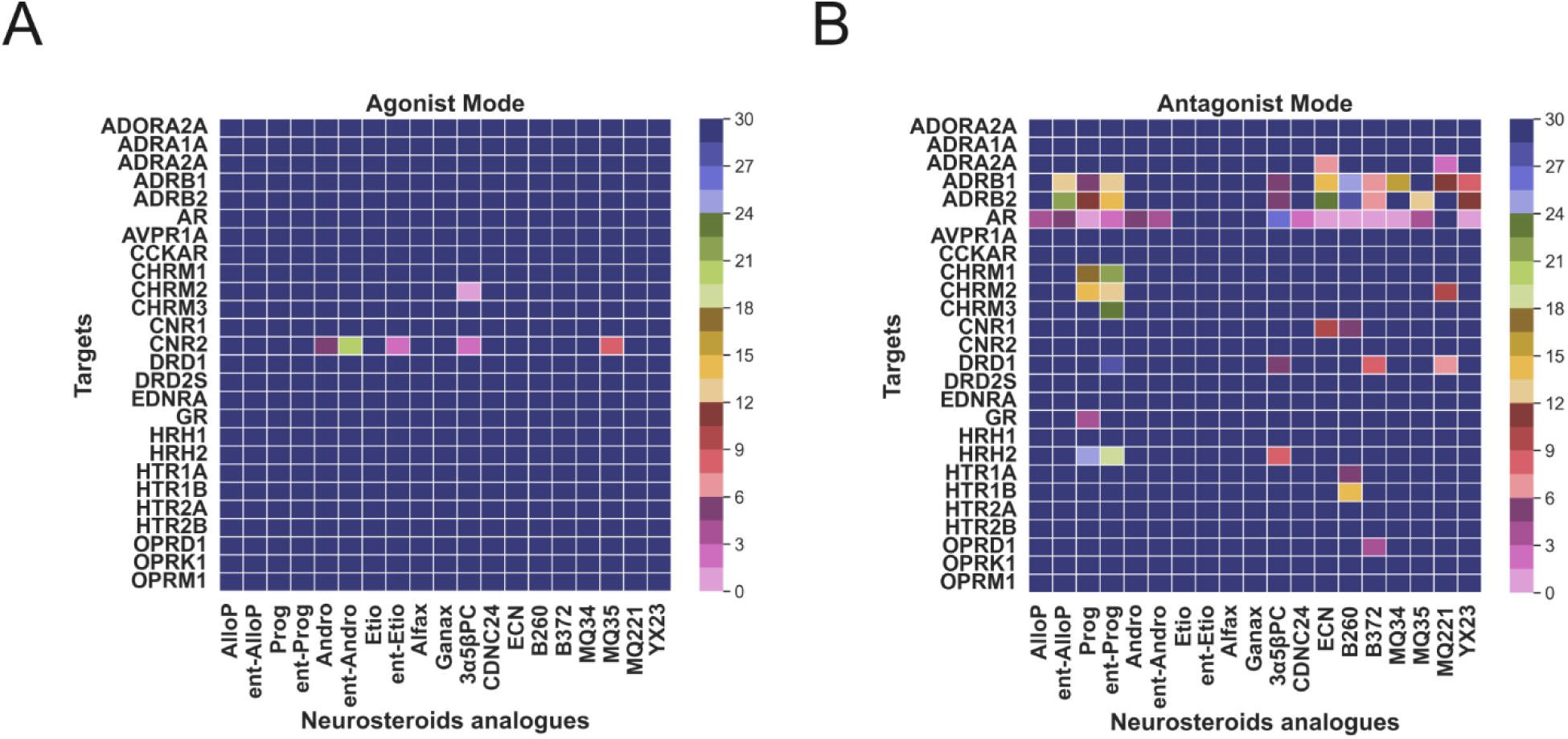
Heatmap of in-vitro activity of neurosteroids and their analogues at various GPCRs and nuclear receptors. **(A)** The heatmap represents the agonist mode activity profiles of a series of neurosteroids and their analogues across multiple GPCRs and nuclear receptors (e.g., AR and GR) **(B)** Heatmap representation of the antagonist mode activity profiles across multiple GPCRs and nuclear receptors (e.g., AR and GR). Scale bar color map transitions from darker shades (representing lower activity) to lighter shades (indicating higher activity). Tick marks are defined at intervals of 3 µM units, ranging from 0 to 30 µM.

### Serotonergic receptor activity

Next, we examined the NS analogues at the serotonin ligand-gated ion channel HRT3A in both blocker and opener modes using the FLIPR Tetra system (Figure 4). The activity dose response curves in Figure S5A demonstrate that eight analogues inhibited HRT3A activity with IC_50_ value ranging from 6.15 µM to 18.21 µM (Table 2). The closely related NS Andro and Etio showed similar inhibitor potency, with IC_50_ values of 7.43 µM and 7.35 µM, respectively. Additionally, the enantiomer of Etio (*ent*-Etio) exhibited similar potency (IC_50_=6.15 µM), whereas *ent*-Andro showed low potency (IC_50_=19.31 µM) compared to Andro (Table 2). Other NS, including ECN (IC_50_=13.25 µm), B260 (IC_50_=18.21 µM), MQ34 (IC_50_=17.55 µM) and the B260 diastereomer (B372), AlloP, and MQ34 showed a clustered IC_50_ range (∼12.33-16.68 µM), somewhat higher than Andro. We also tested serotonin GPCRs including HRT1A and HRT1B (Gαi/o coupled) by cAMP assay and HRT2A and HRT2B (Gαq coupled) by calcium flux assay in agonist and antagonist mode (Figure 5), these receptors are highly involved in various physiological processes including mood and cognition. The concentration-response relationship suggested that only B260 showed antagonistic activity at HRT1A (IC_50_=5.14 µM) and HRT1B (IC_50_=14.58 µM) (Figure S5B & C).

### Adrenergic receptor activity

We evaluated the activity of analogues at adrenergic receptors including ADRA1A, ADRA2A and ADRB1-2 (Figure 5A, B). KEGG pathway enrichment analysis for drug targets suggested that the hsa04261 pathway (adrenergic signaling in cardiomyocytes) was significantly enriched (Figure S3A). This analysis outcome was mirrored in the screening results, most analogues showed activity toward ADRB1-2 (Figure 5B, S6A and B), receptors that are predominantly expressed in cardiomyocytes and regulate cardiac function (Madamanchi, 2007). Prog, 3α5βPC and B372 showed antagonism with IC_50_ values of ∼5-6 µM at ADRB1, good potency compared to other analogues (Table 2, Figure S6A). Weaker antagonism was observed for *ent*-Prog (IC_50_ 13.26 µM), ECN (IC_50_14.58 µM), and YX23 (IC_50_ 8.23 µM), *ent-*AlloP (IC_50_ 13.23 µM), B260 (IC_50_ 25.13 µM), and MQ34 (IC_50_ 15.23 µM) Table 2 (Figure S6A). 3α5βPC and B372 also showed similar antagonistic potency at ADRB2 (4.57 µM and 6.04 µM IC_50_ respectively), while others including MQ35 (IC_50_=12.01 µM), YX23 (10.73 µM), Prog (IC_50_=11.92 µM) and *ent*-Prog (IC_50_=14.26 µM) showed modest antagonistic activity (Figure S6B). Andro, ECN and B260 showed poor potency toward ADRB2 (>20 µM IC_50_). Only ECN (6.82 µM) and MQ221 (1.99 µM) antagonized α adrenergic receptor 2A (ADRA2A) (Table 2; Figure S6C), and none of the steroids showed agonist activity.

### Cannabinoid and other GPCRs receptor activity

Several lines of evidence suggest that retrograde endocannabinoid signaling suppresses GABAergic synapses (Alger, 2002; Barna et al., 2024). Consistent with this, the retrograde endocannabinoid signaling pathway (hsa04723) was identified in our KEGG pathway enrichment analysis, as illustrated in Figure S3A. Despite a few prior studies of NS activity at cannabinoid receptors (CNR) (Vallée et al., 2014; Haney et al., 2023), we assessed the interaction of NS analogues with CNR1 and CNR2 using a cAMP assay.

Interestingly, five compounds exhibited an agonist response with CNR2, while two showed an antagonist response with CNR1 (Figure 5). Specifically, the enantiomer of Etio and 3α5βPC demonstrated CNR2 agonist dose responses with EC_50_ values of 2.86 µM and 2.75 µM, respectively (Figure S7A). Additionally, the endogenous NS Andro exhibited an EC_50_ of 4.67 µM, whereas its enantiomer displayed a substantially lower agonist potency with an EC_50_ value 19.84 µM (Figure S7A). The unnatural enantiomer MQ35 demonstrated a modest agonist response with EC_50_ value 7.82 µM at CNR2, while the natural enantiomer MQ34, exhibited no activity. Only two 5β-reduced compounds, ECN and B260, demonstrated CNR1 antagonism with IC_50_ values of 9.55 µM and 4.89 µM, respectively (Figure S7B).

We tested analogues against several other important families of GPCRs that respond to neurotransmitters/neuromodulators: opioids, histamine, cholecystokinin, and arginine vasopressin (Figure 5). However, with *in vitro* screening, we did not observe notable activity except for the dopamine receptor 1 (DRD1), histamine receptor 2 (HRH2), and delta opioid receptor (OPRD1). Three analogues including 3α5βPC, B372 and MQ221 exhibited DRD1 antagonist activity with similar range of IC_50_ values (5.67-8.25 µM; Figure S7C), while *ent*-Prog had very weak potency (Table 2). Among the tested analogues, only three molecules, including 3α5βPC and Prog, along with its enantiomer, exhibited weak HRH2 antagonistic activity, with IC_50_ values of 8.89 µM, 25.37 µM, and 18.4 µM, respectively (Table 2; Figure S7D). Only B372 exhibited OPRD1 antagonism with 4.25 µM IC_50_ (Figure S7E).

### Ion Channel Activity

Enrichment analysis and beta-adrenergic receptor activity of NS suggested that NS related compounds have some association with regulation of cardiac function. This notion is also supported by several previous studies (Shiyue et al., 2017; Head et al., 2019; Pan et al., 2024). When we examined our enrichment GO analysis (Figure S4), we found ion channel complex (GO:0034702) containing ion channels such as NaV1.5, Ca_v_1.2, KvLQT1 and hERG highly associated with cardiac functions (Grant, 2009). We tested our NS analogues at all four ion channels (Figure 4). Eight of nineteen analogues inhibited Na_v_1.5 channel activity in a dose dependent manner (Figure S8A), in which the enantiomer of Prog (*ent*-Prog) and the carbon 3 diastereomer of B260, B372, have ∼10 µM IC_50_ and others including *ent*-AlloP, Prog, Andro, ECN, MQ34 and MQ35 have IC_50_’s ranging from 16-26 µM, comparatively higher than physiological concentrations of endogenous NS (Table 2). Only B260 exhibited inhibitory activity at Ca_v_1.2, with an IC_50_ of 10.71 µM (Table 2; Figure S8B). All nineteen NS also were tested for KvLQT1 and hERG activity and we did not find any response up to 30 µM (Figure 4). In all enrichment analyses, including GO-term and KEGG, glutamate receptor function was strongly represented (Figure 2B & S3A). Nevertheless, none of the NS showed blocker or opener activity at N-methyl-D-aspartate receptors NMDAR (1A/2B) (Figure 4A and B). We note that our compound list contains at least two NS with previously documented inhibitory activity at neuronal NMDARs (Zhu and Vicini, 1997; Mennerick et al., 2001). We address caveats of the screening assays in the Discussion.

### Neurotransmitter metabolic enzymes & transporters

Given actions of steroids at some neurotransmitter metabolic enzymes, we tested all nineteen NS analogues at AChE and MAOA. Only Prog was active, inhibiting AChE with IC_50_ value 12.48 µM (Figure 6 and S10C). Neurotransmitter transporters are essential for regulating the concentration of neurotransmitters in the synaptic cleft and ensuring the precise control of neurotransmitter signaling (Torres et al., 2003). We tested NS analogues activity at multiple neurotransmitter transporters including SERT, DAT, and NET. None of the compounds had effects on any of the transporters (Figure 6).

**Figure 6.**
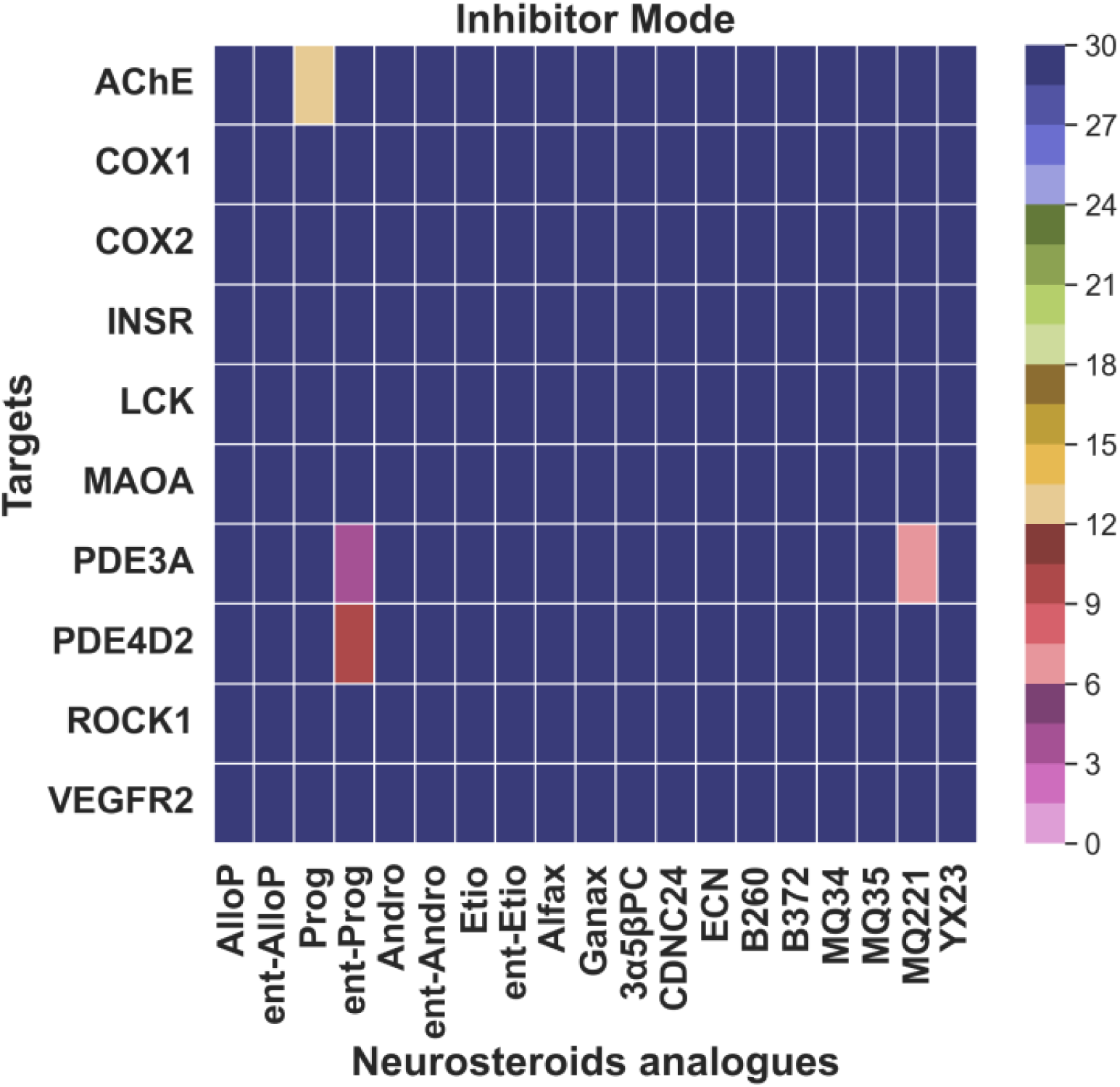
Heatmap of enzymatic activity of neurosteroids and their analogues at various enzymes. The heatmap represents the activity profiles of a series of neurosteroids and their analogues across multiple enzymatic activities, including kinase in inhibitor mode. Scale bar color map transitions from darker shades (representing lower activity) to lighter shades (indicating higher activity). Tick marks are defined at intervals of 3 µM units, ranging from 0 to 30 µM.

### Nuclear hormone receptor activity

Unsurprisingly, network pharmacology drug-target enrichment data, along with previous work, showed that steroid-like molecules are associated with the GO-term GO:0003707 (nuclear steroid receptor activity; Figure 2B). We focused on nuclear receptor GR and AR for our investigation of NS analogues molecules activity because of limited exploration with NS analogues. Understanding the direct activity on GR and AR is crucial for elucidating their potential effects on various physiological processes. We investigated the activity of the 19 analogues at GR and AR using nuclear protein interaction and nuclear translocation assays, respectively in agonist and antagonist mode (Figure 5). Dose response results suggested that only Prog showed GR antagonist activity with an IC_50_ value of 3.61 µM (Table 2; Figure S10B). Concentration response of AR activity suggested that all compounds antagonized AR activity except sulphated compound MQ221, and both enantiomers of Etio (Figure S10A), which highlights the differential activity of NS and their analogues at androgen receptors. Notably, the molecules B260, B372, ECN, MQ34, Prog, and YX23 exhibited the most potent antagonistic activity, with IC_50_ values of 0.53 µM, 0.75 µM 0.16 µM, 0.12 µM, 0.03 µM and 0.60 µM respectively (Table 2). Interestingly, the unnatural enantiomers of MQ34 (i.e., MQ35) and Prog (*ent*-Prog) exhibited IC_50_ values of 4.27 µM and 2.92 µM respectively; the natural enantiomers were ∼35-100 times more potent (Table 2). Other analogues such as CDNC24, AlloP, and Andro also demonstrated moderate antagonistic effects, with IC_50_ values ranging from 2.52 µM to 4.82 µM (Table 2). Additionally, unnatural enantiomers of AlloP and Andro (e.g., *ent*-AlloP and *ent*-Andro) displayed activity similar to the natural molecule, with IC_50_ values ranging from 4.12 µM to 4.82 µM. However, 3α5βPC exhibited weaker antagonistic activity at AR, with an IC_50_ value of 26.95 µM.

### Kinases and non-kinase enzymes

Some NS-like molecules have shown association with kinase signaling activity, as indicated by our KEGG pathway analysis and previously published work (Adams et al., 2015). Nineteen analogues were evaluated for kinase activity at LCK, VEGFR2, ROCK1, and INSR; no changes were observed (Figure 6). Additionally, we assessed AlloP and Prog, along with their enantiomers at single concentration (10 µM), for their kinase inhibition activity at 481 kinases. However, we did not observe notable kinase activity inhibition by these NS (Table S3). Furthermore, we investigated the inhibitory effects of all NS on phosphodiesterases (PDEs) and cyclooxygenases (Cox). The results are illustrated in Figure 6. Interestingly, the enantiomer of Prog inhibited the activity of PDE3A and PDE4D2 in a dose dependent manner with IC_50_ values of 3.45 µM and 9.76 µM, respectively while sulphated NS MQ221 inhibited PDE3A response with 7.05 µM IC_50_ (Table 2; Figure S9 A and B). None of the NS analogues showed inhibitory activity against Cox-1 and Cox-2 enzymes.

## DISCUSSION

To date, the most explored mechanism of action for NS analogues is modulation of GABA-A receptors. Previously, several reports of our research group and others extensively investigated GABA-A receptor activity of NS, including natural and synthetic analogues. Several other reports, however, suggest that multiple other targets are modulated by NS (Darbandi-Tonkabon et al., 2003; Cheng et al., 2019; Covey et al., 2023; Balan et al., 2024). Nevertheless, we still lack a collective understanding about multi-target activity of NS. Here, we broadly characterized the activity of NS analogues, including natural and synthetic analogues and enantiomeric compounds at several therapeutically important but under-explored targets. The compounds evaluated included pregnane steroids and polar/charged compounds, along with androstanes. We applied a computational approach to uncover classes of targets and followed with *in vitro* screening to experimentally explore specific targets with selected NS analogues from different structural classes. Interestingly, our results suggest that natural and synthetic NS analogues show multi-target activity, some with relatively high potency.

Our computational analysis suggested that several potential therapeutic molecules with structural similarity to AlloP bind to nAChRs, consistent with published work (Paradiso et al., 2000, 2001; Steinbach and Akk, 2011). More than half of our tested NS analogues blocked the nAChR α4β2 receptor and some of these exhibited IC_50_ values less than 5 µM. Nicotinic receptors are crucial for cognitive function, addiction mechanisms, and mood regulation, making these receptors a significant therapeutic target. NS block of α4β2 nAChRs could reduce nicotine’s rewarding effects, supporting smoking cessation efforts. Additionally, the dual modulatory properties of NS, affecting both GABA-A receptors and nAChRs, could provide a novel strategy for managing mood disorders and pain. Interestingly, 3α5βPC functions as a strong CHRM2 agonist without nAChR activity, and two others are weak CHRM2 antagonists. The muscarinic activity of 3α5βPC appears to complement effects at GABA-A receptors and NMDA receptors (Mennerick et al., 2001). These ionotropic receptor effects did not appear among screening results, possibly because of potency but also raising the possibility of false negative results in the screens.

Previous studies suggested that several NS including Prog and AlloP functionally antagonize serotonin HRT3 receptors (Barann et al., 1999; Rupprecht and Holsboer, 2001). Our results suggest that other NS also block HRT3A ionotropic receptors. At serotonin GPCRs, only B260 was active and antagonized HRT1 A and B receptors. Among other functions, these receptors modulate neurotransmitter release, including serotonin and dopamine. Also, antidepressant medications may modulate the level of endogenous NS (Broekhoven et al., 1998; Serra et al., 2001; Broekhoven and Van, 2003; Veska and Sampson, 2006). This bidirectional interaction highlights the complex relationship between NS and the serotonergic system, an interplay that could be harnessed for therapeutic benefit.

Chronic hypertension enhances NS levels and AlloP has shown antihypertensive effects in mice with neurogenic hypertension (Stevenson et al., 2017; Head et al., 2019). Consistent with this, our results showed AlloP antagonism of adrenergic receptors, including ADRB1, ADRB2, and ADRA, which are predominantly expressed in cardiomyocytes. This antagonism suggests a potential modulation of the sympathetic nervous system by NS, which could impact a range of physiological responses such as heart rate, blood pressure, and stress-related behaviors. Correlation studies have shown that elevated levels of certain NS are associated with reduced adrenergic activity, highlighting a possible inverse relationship between NS concentration and adrenergic receptor activation (Gunn et al., 2004, 2015; Morita et al., 2004). For example, in conditions characterized by hyperactivity of the adrenergic system, such as hypertension or anxiety disorders, increased levels of these NS could serve as a natural counterbalance to excessive adrenergic stimulation. Thus, NS could help maintain homeostasis in the autonomic nervous system. At the Ca_v_1.2 calcium channel, which is important in cardiovascular physiology, only B260 exhibited blocking activity. B260 has also been studied as a blocker as low voltage activated Ca^2+^ channels (Joksimovic et al., 2020). Importantly, NS were largely inert at KvLQT1/minK and hERG, key cardiac liability targets.

NS also blocked the activity of voltage dependent sodium channel (Na_v_1.5) with moderate to high IC_50_ values. This mechanism offers another potential point at which NS can inhibit electrical activity of neurons, with therapeutic potential in conditions like epilepsy and mood disorders. Along with adrenergic and cholinergic GPCRs, we tested other GPCRs including CNR1-2, DRD1-2, HRH1-2, EDNRA and opioid receptors that play crucial roles in normal physiology. Several lines of evidence suggest that retrograde endocannabinoid signaling suppresses GABAergic synapses (Ohno-Shosaku et al., 2001; Alger, 2002), but limited previous evidence suggests direct involvement of NS with cannabinoid signaling (Vallée et al., 2014; Raux et al., 2022; Raux and Vallée, 2023). Interestingly, five molecules exhibited agonist activity at CNR2, while two NS showed an antagonist response with CNR1. Thus, we confirm an interaction with cannabinoid receptors, with implications for reward modulation (Romieu et al., 2003; Seib et al., 2023). DRD1 and OPRD1 are also both highly involved in reward-related behavior (Merrer et al., 2009; Seib et al., 2023). We found that B372 antagonizes both receptors with reasonable IC_50_, while 3α5βPC and MQ34 antagonized the DRD1 receptor only.

Growing evidence suggests that NS analogues modulate enzyme activity including kinases. We tested NS for enzyme activity including AChE, COX1-2, INSR, MAOA, phosphodiesterase’s and a panel of kinase enzymes. MQ221 inhibited the phosphodiesterase PDE3A that is highly involved in cardiovascular functions. The enantiomer of Prog inhibited both PDE3A and PDE4D2. Inhibition of PDE4D2 and PDE3A by the enantiomer of Prog suggests broader effects on cyclic nucleotide signaling pathways beyond cardiovascular functions. These enzymes play multiple roles in physiological processes, including neuronal signaling, inflammation, and immune responses (Michael-Claude et al., 2007; Omori and Kotera, 2007). Some studies suggest that Prog can affect the activity of AChE in certain experimental paradigms (Lakomy et al., 1986; Amin et al., 2020); our result also found the same trend with a moderate IC_50_ value. Further research is required to fully understand the mechanisms and the physiological and clinical implications of AChE inhibition by Prog.

In general, NS are inert to nuclear hormone receptor activity, but the screening results suggest that some NS also modulate activity of nuclear steroid hormone receptors. Steroid hormones bind cognate receptors at nanomolar concentrations; the micromolar concentrations identified herein represent comparatively low affinity. Nevertheless, most of the NS analogues blocked AR activity. Interestingly, only Prog exhibited GR inhibition; none exhibited activation. This suggests a prevalent capacity among NS to inhibit androgenic signaling pathways in the brain and potentially other tissues. These findings underscore the potential of NS to modulate androgen-driven processes in physiological and pathological states. Therapeutic implications of this AR antagonism are substantial, particularly in the context of neuropsychiatric and neurodegenerative disorders. By blocking AR, NS may mitigate the neurotoxic effects of excessive androgen activity, offering neuroprotective benefits in conditions such as Alzheimer’s disease and traumatic brain injury. These findings pave the way for the development of NS based therapies, leveraging their AR antagonistic properties to address a range of conditions associated with aberrant androgen signaling.

## LIMITATIONS OF THE STUDY

We identified several limitations, likely not comprehensive, in the virtual screens. Some important neurosteroids, such as the canonical sulfated NS pregenolone sulfate (Majewska and Schwartz, 1987; Wu et al., 1991), are not represented in the Drug Repurposing Hub. The Hub also does not cover some known mechanisms of action, such as the GABAergic actions of 17-PA, a compound that our group synthesized (Mennerick et al., 2004). Furthermore, it identified a mechanism of action for 17-PA (glucocorticoid receptors) that we could find no literature support for.

*In vitro* screening also presents several limitations that should be carefully considered, including the possibility of both false positives and false negatives. Validation of the present hypothesis-generating screening results will be important. Screening systems may not accurately represent the native expression patterns, subunit diversity, and signaling systems found in neurons. Also, the specific conditions of the assay or choice of output measure could affect results. As a result, NS that show little to no activity in these screens might still interact with targets in a physiological setting, leading to potential underestimation of their modulatory effects. For instance, ganaxalone has been studied in preclinical and clinical studies but was surprisingly inert in the present studies. False or misleading positives are also a concern. For instance, numerous NS that have been repeatedly shown to possess GABA-A receptor PAM activity showed GABA-A opener activity but also blocker activity in these screens (Table 2). The basis for the blocker activity is unclear but consistent with effects of high steroid concentrations (Hauser et al., 1996; Foll et al., 1997; Zhu and Vicini, 1997) and with site identification work (Sugasawa et al., 2020). Although direct gating (opening) is also a typical effect of GABA-A receptor PAMs at high concentrations, neither opener nor blocker activity is likely to relate directly to PAM effects. In summary, our work reinforces the pleiotropic signaling pathways of NS analogues and suggests that different combinations of effects can be leveraged by different analogues to produce desired therapeutic outcomes. Although structural elements did not map neatly onto physiological targets, further work with additional analogues could reveal clearer structure-activity relationships. Finally, initial screening outcomes here should be validated in relevant tissue and organ systems to ensure relevance.

## Supporting information

Activity Data as Table S3

## Data availability

The data that support the findings of this study are available from the corresponding author upon reasonable request.

## ACKNOWLEDGEMENTS

This work was supported in part by R01 MH123748 (SM) and P50 MH122379 (SM and CFZ).

## AUTHOR CONTRIBUTIONS

DFC, SM, AB, and CFZ designed the screening experiments. AK designed and performed computational analyses and wrote a first draft. All the authors participated in analysis and editing the manuscript.

## DELCLARATION OF GENERATIVE AI AND AI-ASSISTED TECHNOLOGIES IN THE WRITING PROCESS

During the preparation of this work the author(s) used ChatGPT and Microsoft Word grammar suggestions to ensure grammatical and stylistic flow. After using these tools, the authors reviewed and edited the content as needed and take full responsibility for the content of the publication.

## SUPPLEMENTARY INFORMATION

**Figure S1.**
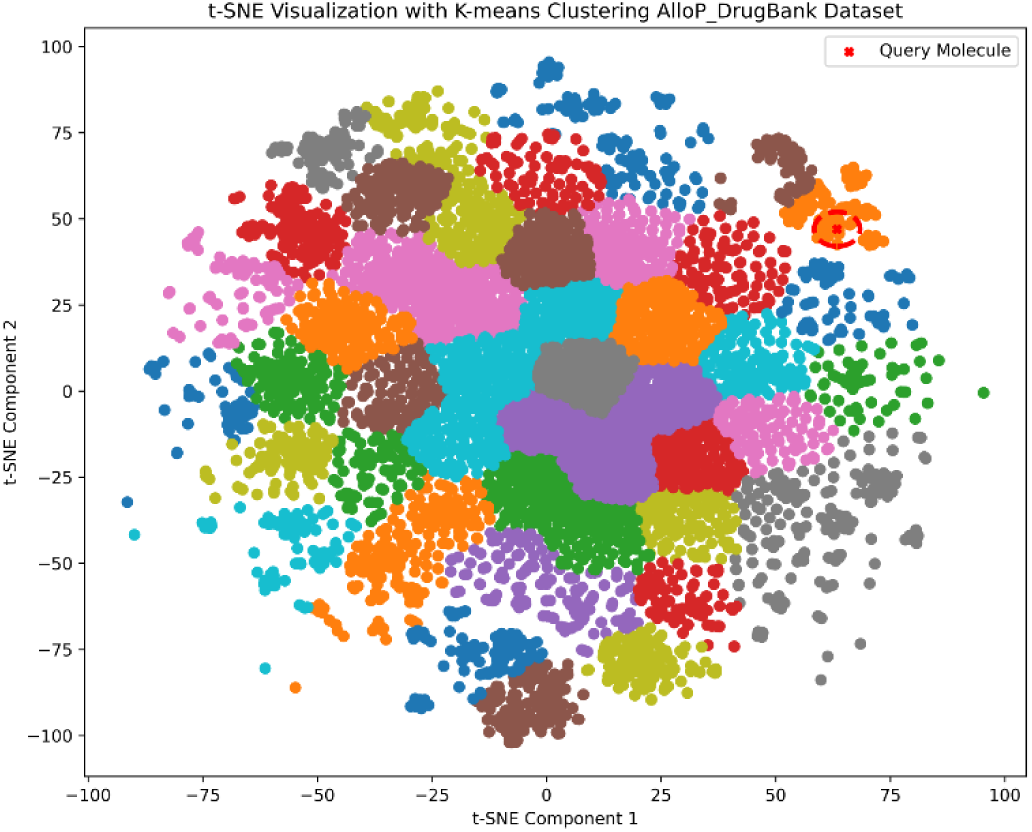
t-SNE based chemical similarity analysis of AlloP with the DrugBank molecule dataset. Figure illustrates t-SNE visualization of DrugBank molecule dataset with query molecule (AlloP) in 2D chemical space. The analysis was performed using 4096-bit Morgan fingerprinting with default t-SNE setting for embedding. Molecules similarity determined by k-nearest neighbors’ analysis and chemical space distance from AlloP, a circle with a radius of 5 units was plotted around the query molecule in the t-SNE visualization.

**Figure S2.**
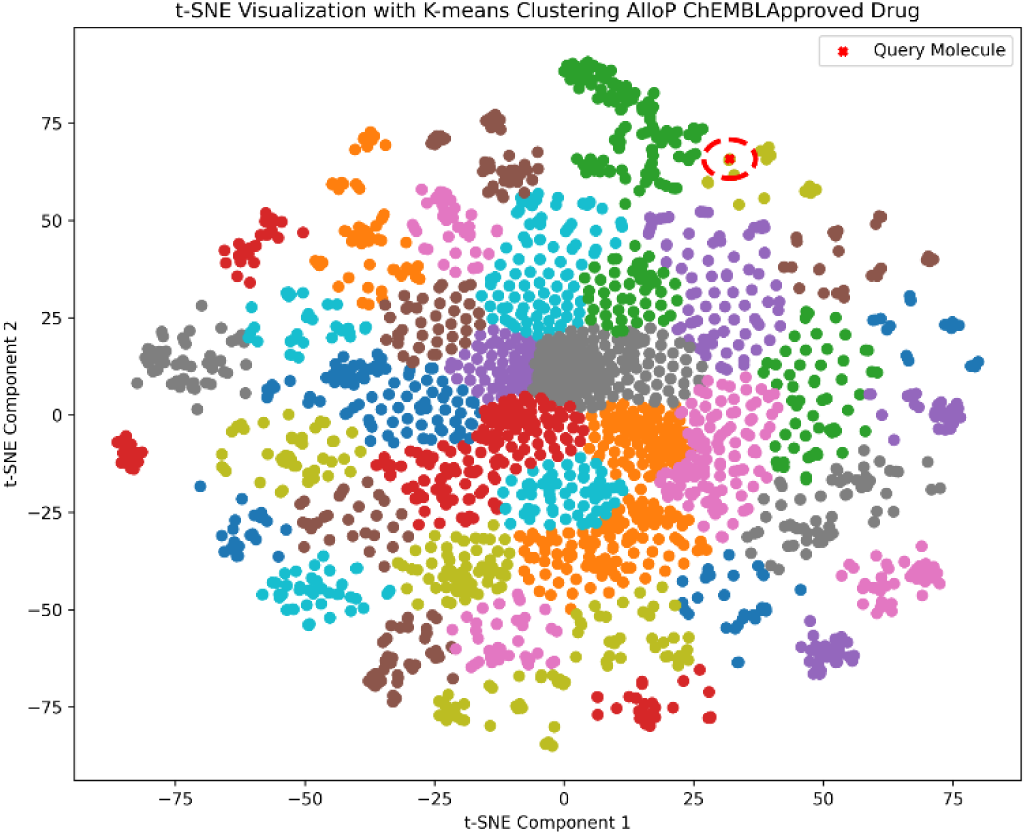
t-SNE visualization of ChEMBL approved drug molecule dataset with query molecule (AlloP) in 2D chemical space to determine neighbor molecule from approved drug dataset.

**Figure S3.**
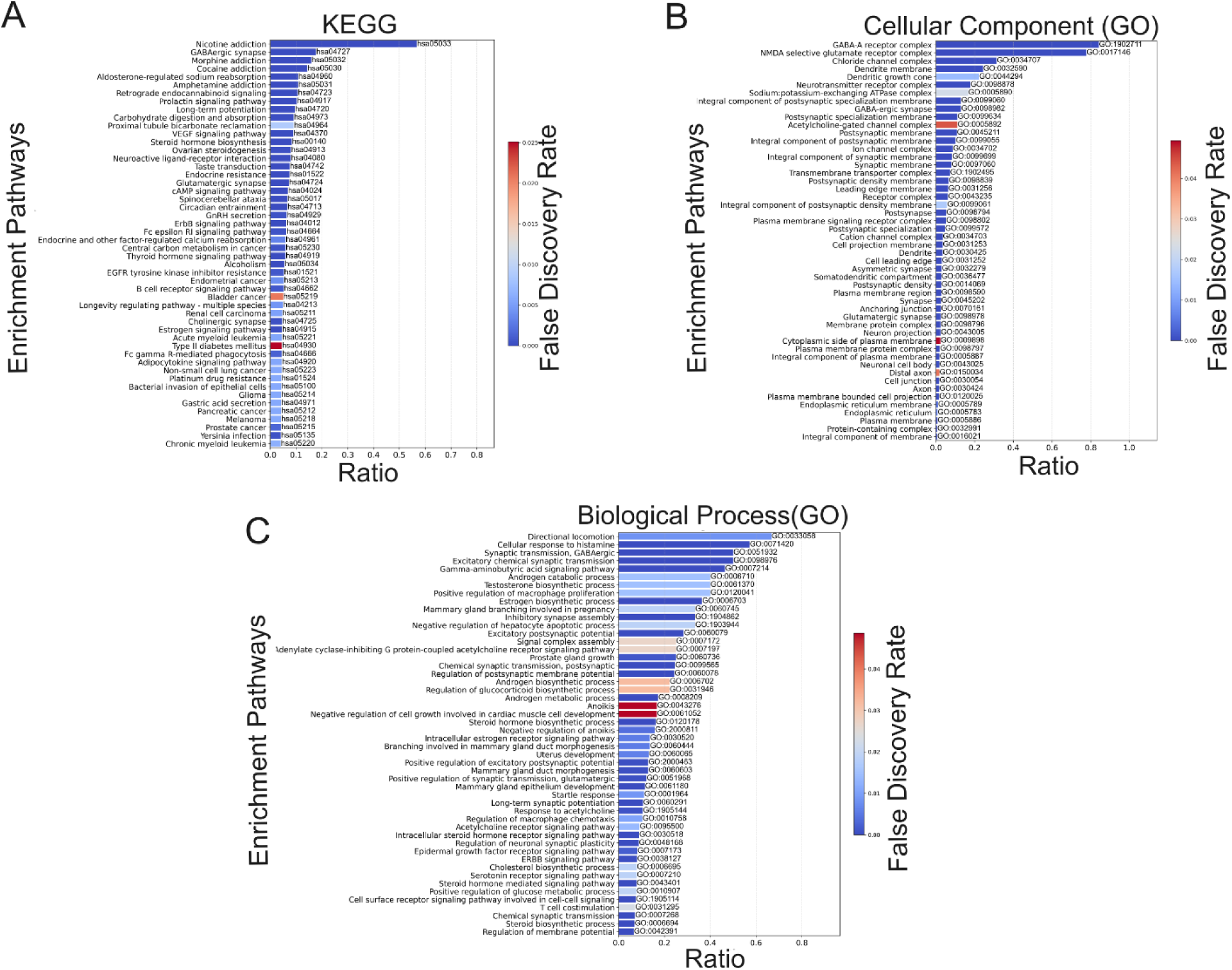
KEGG Pathway and Gene Ontology (GO) enrichment analysis. **(A)** The graph illustrates subsequent KEGG pathway gene enrichment analysis associated with AlloP and neighbor molecule target genes. **(B)** Gene ontology (GO) enrichment analysis of the network identified significantly overrepresented cellular component associated with the drug target genes. **(C)** List of enriched pathway gene associated with biological process. These enriched functions are depicted in the associated bar chart, with each bar representing a distinct category, ranked by significance false discovery rate and ratio of the number of genes associated with a particular pathway to the total number of genes.

**Figure S4.**
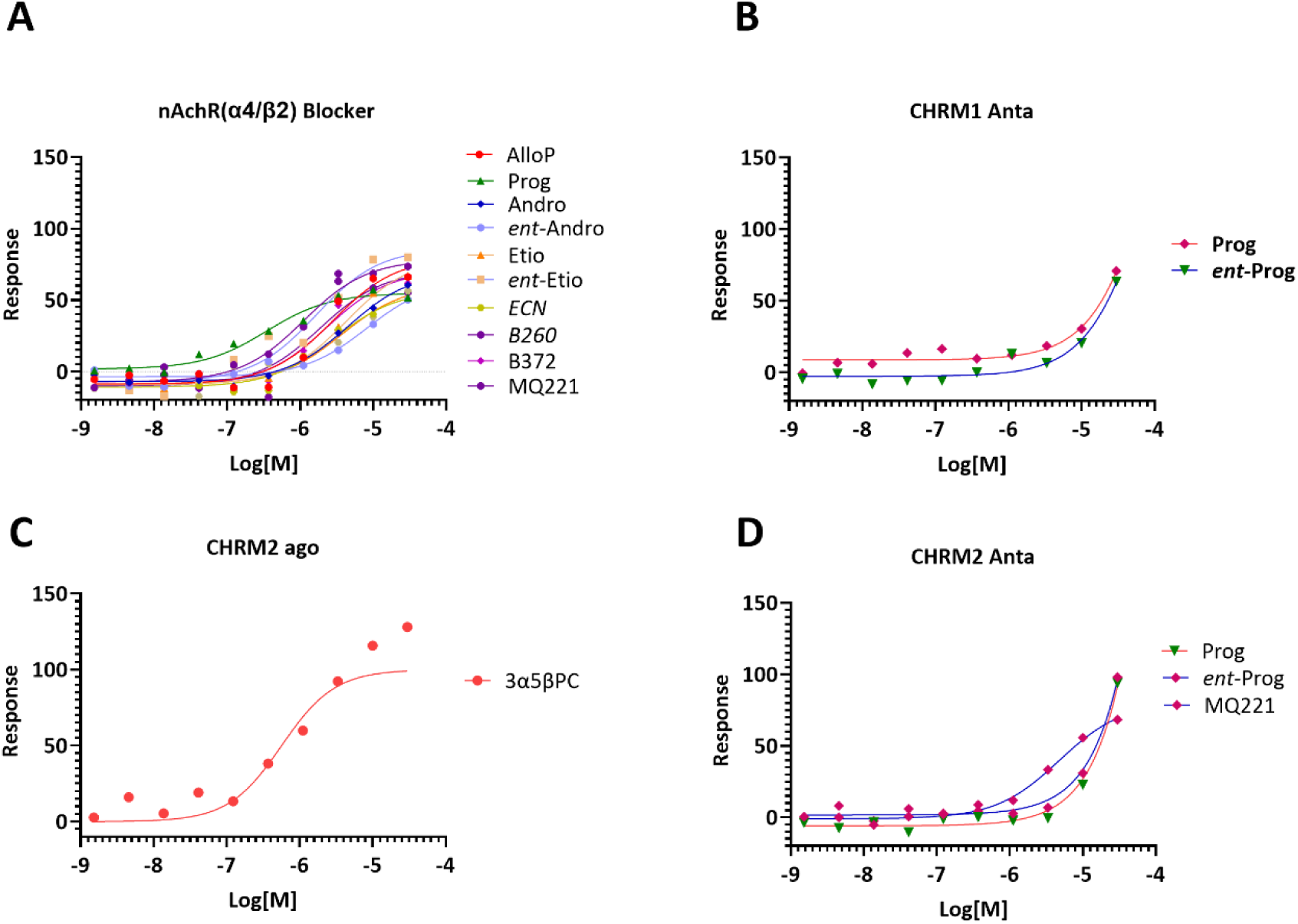
Concentration dependent response activity of NS molecules at cholinergic receptors. (A) Dose response curve for inhibition of nAChR (α4β2) ion channel activity, (B) antagonist activity of CHRM1 receptor, (C) Agonist activity of CHRM2, and (D) Antagonist activity of the CHRM2 receptor.

**Figure S5.**
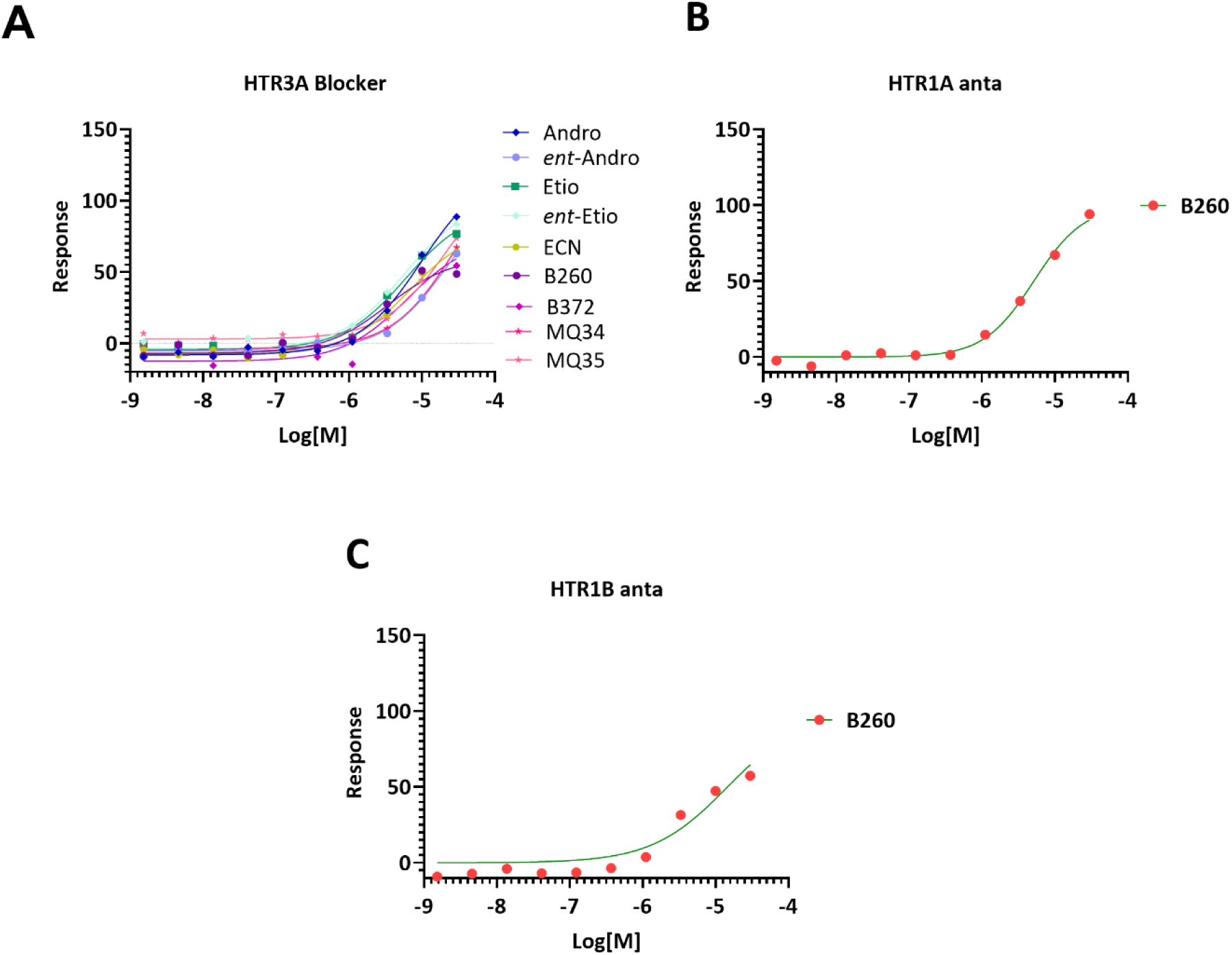
Concentration dependent responses of NS molecules at serotonergic receptors. (A) illustrates inhibition of HTR3A ion channel activity, (B) illustrates antagonist activity of HTR1A receptor, and (C) illustrates antagonist activity of HTR1B receptor

**Figure S6.**
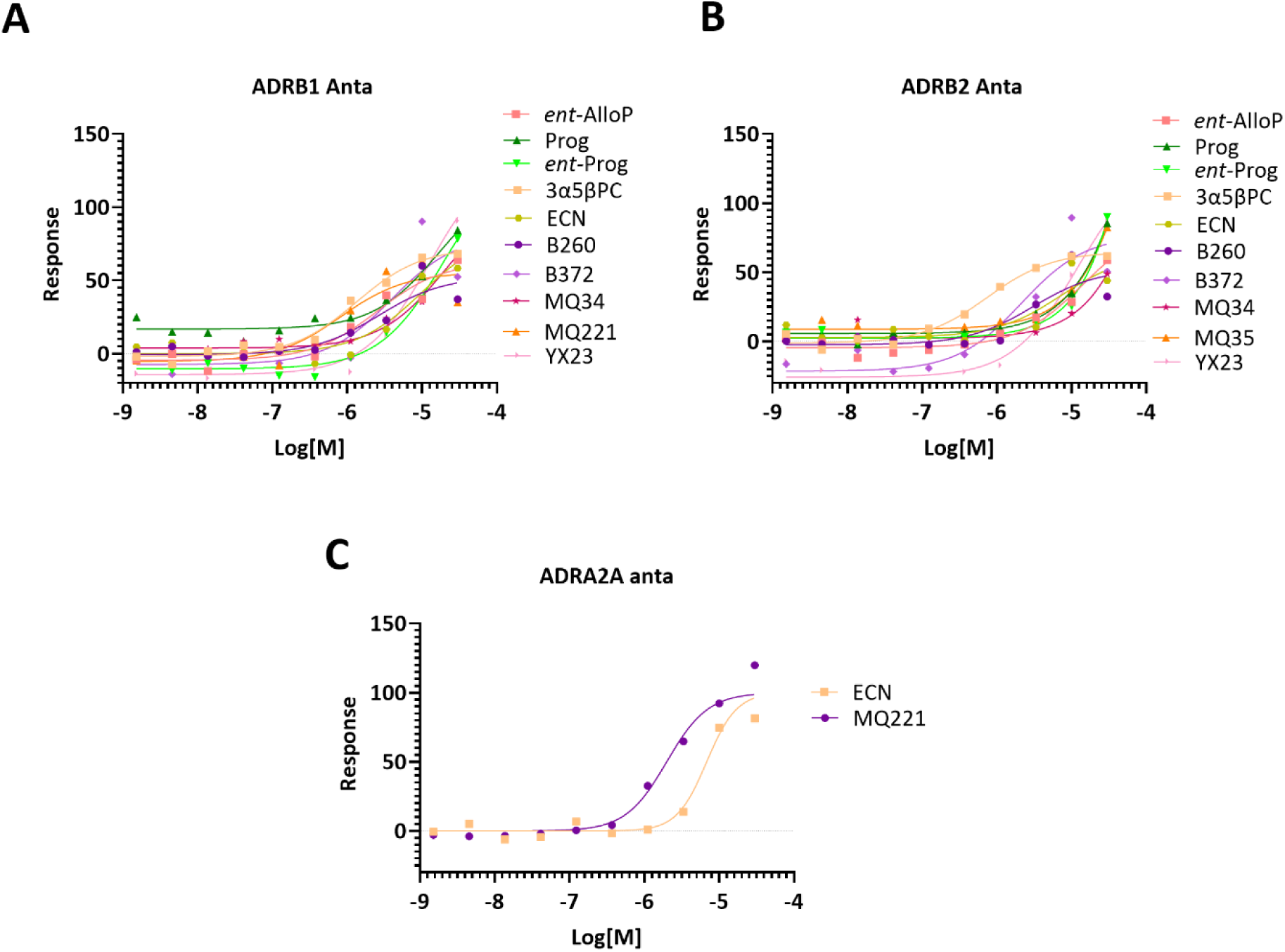
Concentration dependent response of NS molecules: (A) illustrates antagonist activity of ADRB1 receptor, (B) illustrates antagonist activity of ADRB2 receptor, and (C) illustrates antagonist activity of ADRA2A receptor.

**Figure S7.**
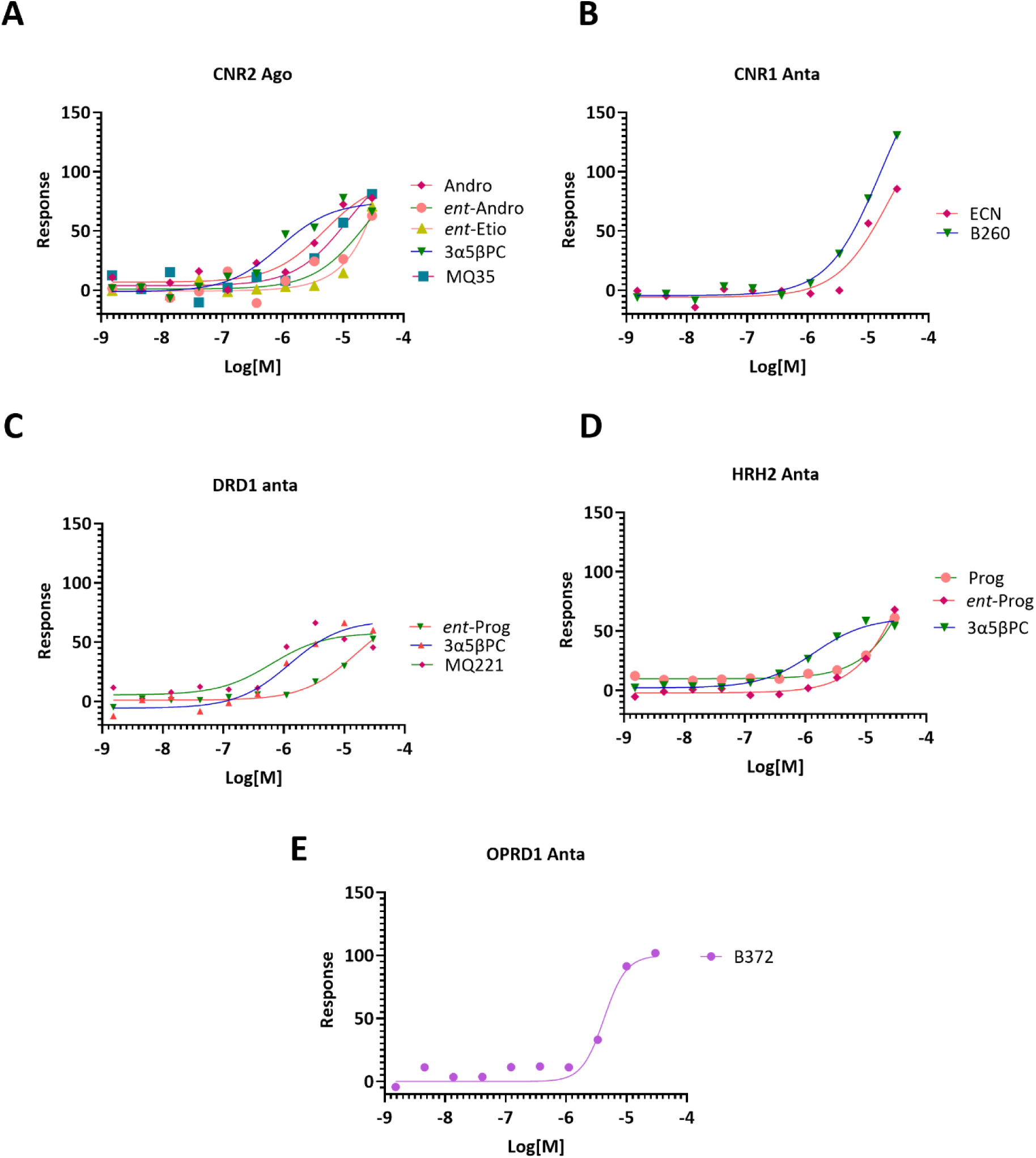
Concentration dependent response of NS molecules: (A) shows agonist activity of CNR2 receptor, (B) illustrates antagonist activity of CNR1 receptor, (C) illustrates antagonist activity of DRD1 receptor, (D) illustrates antagonist activity of HRH2 receptor, and (E) illustrates antagonist activity of OPRD1 receptor.

**Figure S8.**
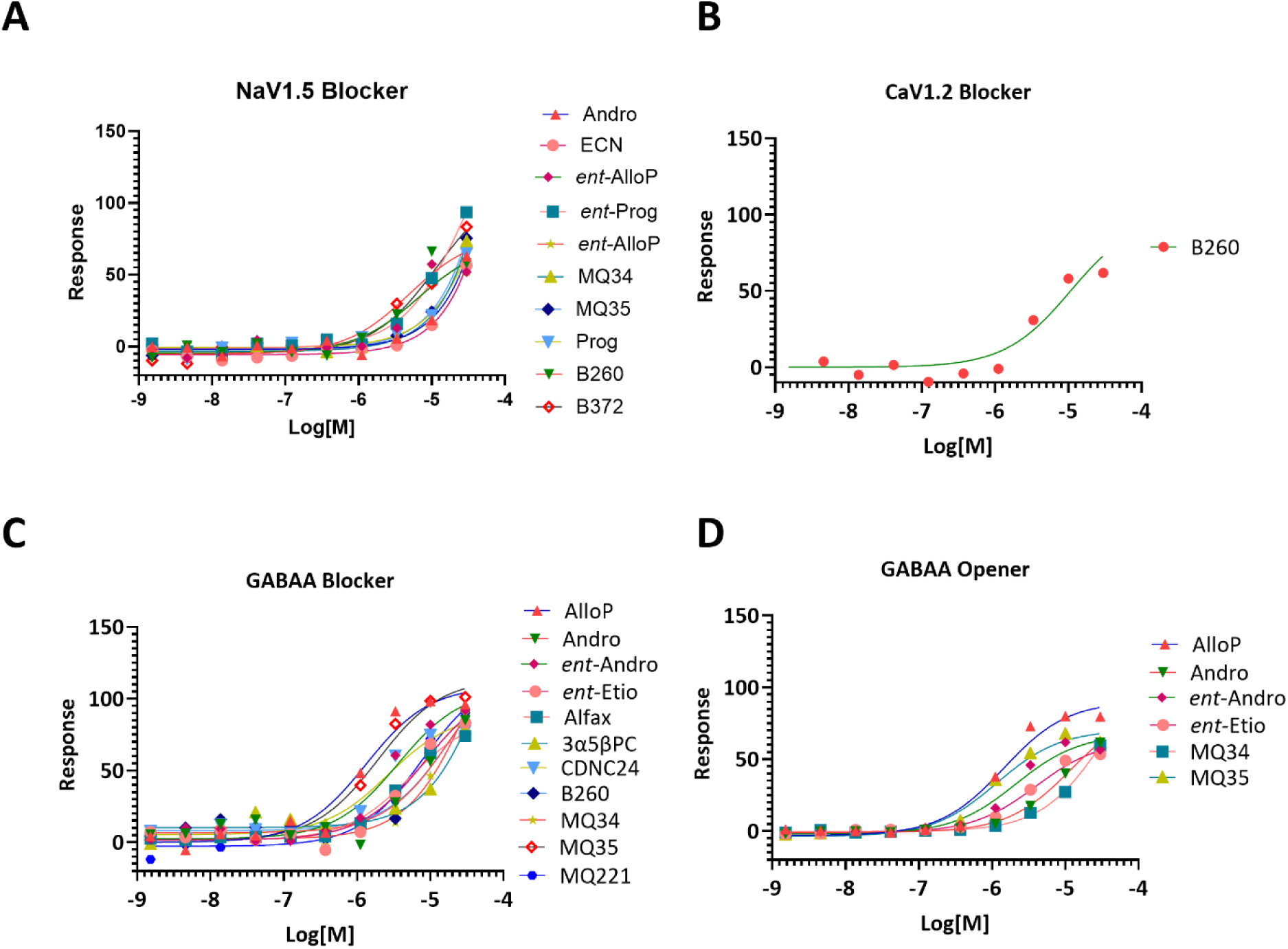
Concentration dependent response of NS molecules: (A) illustrates inhibition of NaV1.5 ion channel activity, (B) illustrates inhibition of CaV1.2 ion channel activity, (C) shows inhibition of GABAA ion channel activity in blocker mode, and figure (D) shows GABAA ion channel activity in opener mode

**Figure S9.**
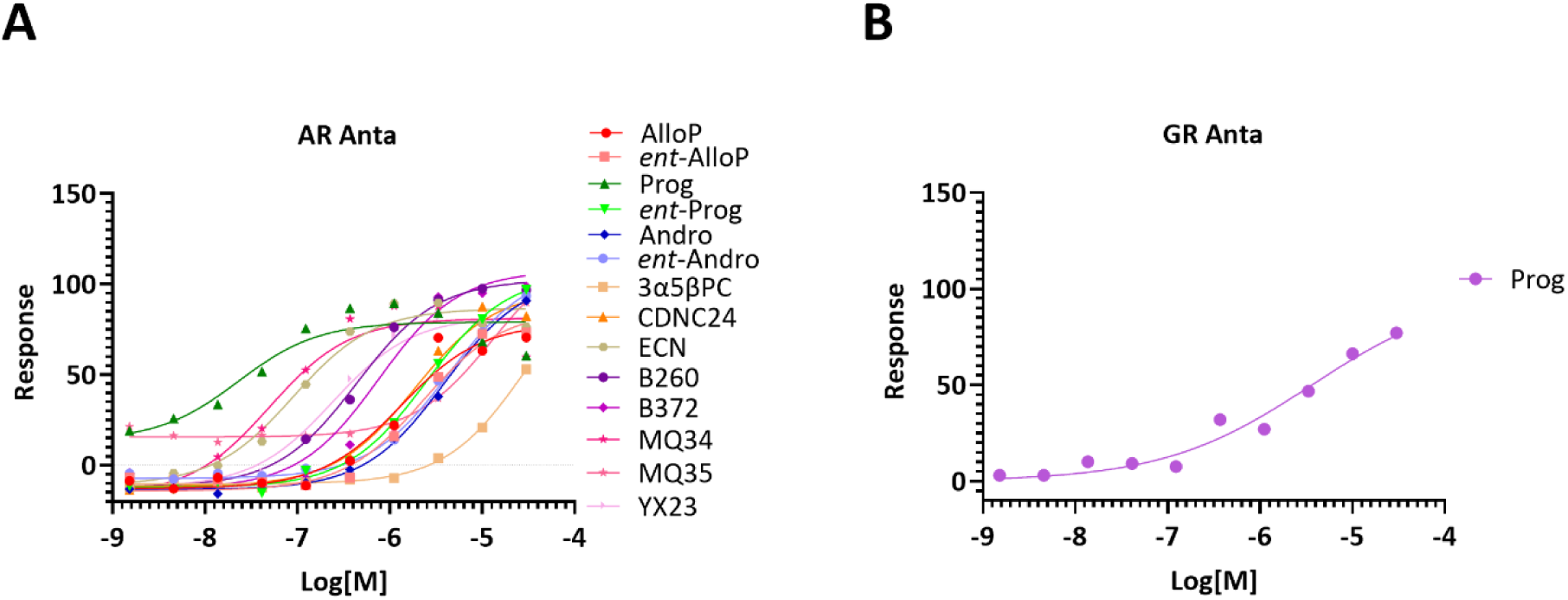
Concentration dependent response of NS molecules at nuclear receptor: (A) illustrates inhibition of AR receptor activity, and (B) shows inhibition of GR receptor activity.

**Figure S10.**
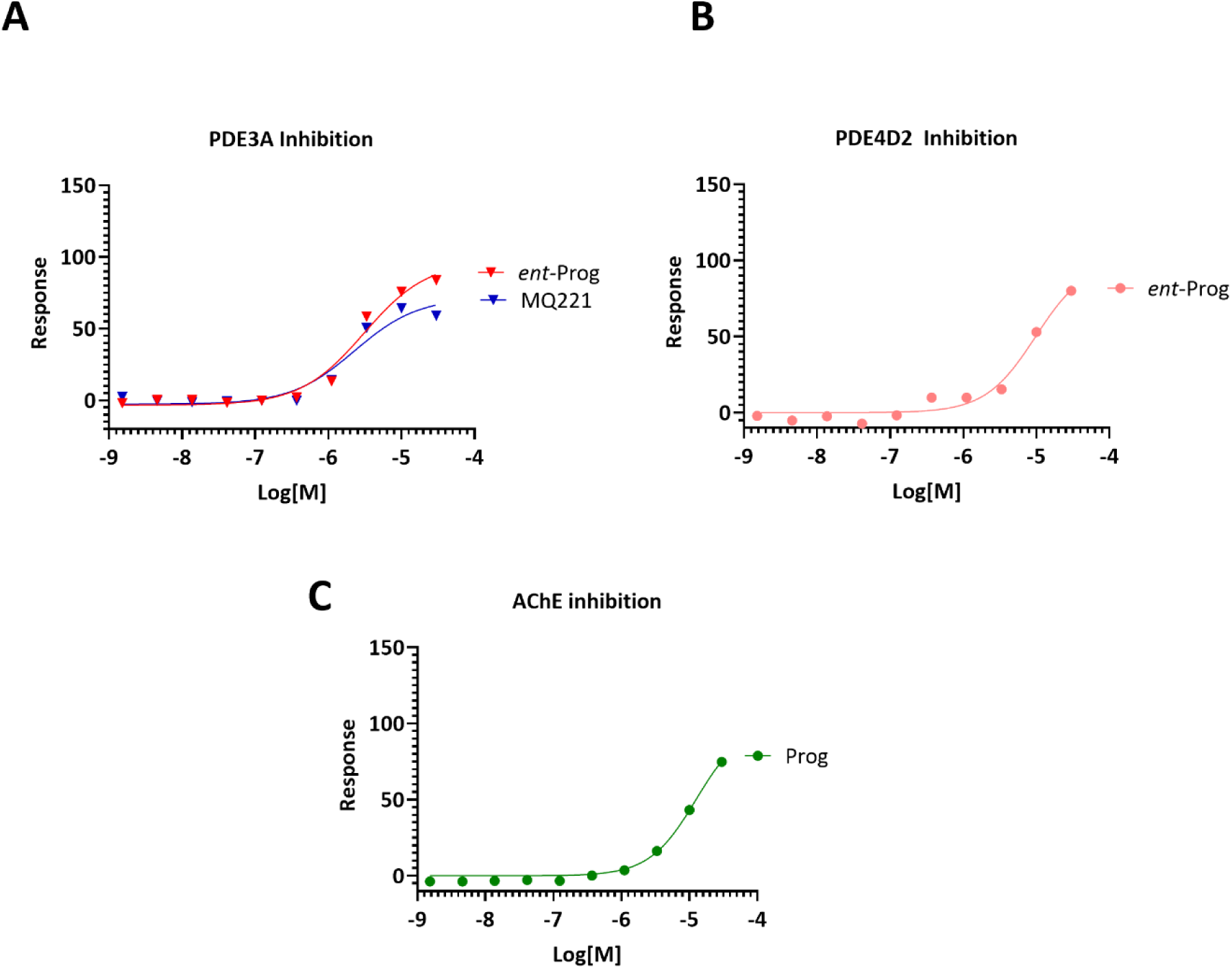
Concentration dependent response of NS molecules at enzyme activity inhibition: (A) illustrates inhibition of PDE3A enzyme activity, and figure (B) shows inhibition of PDE4D2 enzyme activity, and figure (C) shows inhibition of AChE enzyme activity.

