## Supplemental tables for "Multifaceted Actions of Neurosteroids"

**Table S1.** Neighboring molecules and reported mechanisms of action from DrugBank dataset

| DrugBank ID | Common name | Mode of action/Target Name |
|---|---|---|
| DB00621 | Oxandrolone | Androgen receptor |
| DB00728 | Rocuronium | Neuronal acetylcholine receptor subunit alpha-2, Muscarinic acetylcholine receptor M2, 5-hydroxytryptamine receptor 3A |
| DB00858 | Drostanolone | Androgen receptor |
| DB01108 | Trilostane | 3 beta-hydroxysteroid dehydrogenase/Delta 5-->4-isomerase type 1, 3 beta-hydroxysteroid dehydrogenase/Delta 5-->4-isomerase type 2, Estrogen receptor alpha, Estrogen receptor beta |
| DB01216 | Finasteride | 3-oxo-5-alpha-steroid 4-dehydrogenase 2, 3-oxo-5-alpha-steroid 4-dehydrogenase 1, 3-oxo-5-beta-steroid 4-dehydrogenase |
| DB01337 | Pancuronium | Neuronal acetylcholine receptor subunit alpha-2, Muscarinic acetylcholine receptor M2, Muscarinic acetylcholine receptor M3 |
| DB01338 | Pipecuronium | Neuronal acetylcholine receptor subunit alpha-2, Muscarinic acetylcholine receptor M2, Muscarinic acetylcholine receptor M3 |
| DB01339 | Vecuronium | Neuronal acetylcholine receptor subunit alpha-2 |
| DB01451 | 1-Androstenedione | Steroids |
| DB01479 | BA-2664 | |
| DB01481 | 1-Testosterone | Androgen receptor |
| DB01503 | 1-Androstenediol | |
| DB01513 | 17α-methyl-3β,17β-dihydroxy-5α-androstane | |
| DB01514 | Furazabol | |
| DB01530 | 5alpha-androstane-3alpha,17beta-diol | |
| DB01561 | Androstanedione | Estradiol 17-beta-dehydrogenase 1, Steroid Delta-isomerase |
| DB01572 | Methyl-1-testosterone | |
| DB01889 | 16,17-Androstene-3-Ol | Nuclear receptor subfamily 1 group I member 3 |
| DB02854 | Aetiocholanolone | Ig gamma-2 chain C region, Bile salt sulfotransferase, 3-alpha-(or 20-beta)-hydroxysteroid dehydrogenase, Estradiol 17-beta-dehydrogenase 11 |
| DB02901 | Stanolone | Estrogen receptor alpha, Mineralocorticoid receptor, Androgen receptor, Estradiol 17-beta-dehydrogenase 1 |
| DB03882 | 5-Alpha-Androstane-3-Beta,17beta-Diol | Estrogen receptor beta |
| DB03926 | 5alpha-androstane-3beta,17alpha-diol | |
| DB04834 | Rapacuronium | Muscarinic acetylcholine receptor M2 |
| DB05087 | Ganaxolone | GABA(A) Receptor |
| DB05107 | 16-Bromoepiandrosterone | Glucose-6-phosphate 1-dehydrogenase |
| DB05263 | Caprospinol | |
| DB05450 | Smilagenin | |
| DB06157 | Istaroxime | Sarcoplasmic/endoplasmic reticulum calcium ATPase 2, Sodium/potassium-transporting ATPase subunit alpha-1 |
| DB06307 | Apoptone | Apoptosis regulator Bcl-2 |
| DB06412 | Oxymetholone | Androgen receptor, Natriuretic peptides B |
| DB06622 | 7-beta-Hydroxyepiandrosterone | |
| DB06718 | Stanozolol | Androgen receptor, Glucocorticoid binding proteins |
| DB07375 | Etiocholanedione | Ig gamma-2 chain C region, Ig gamma-1 chain C region, Ig kappa chain C region, Ig kappa chain V-II region RPMI |
| DB07447 | 5beta-dihydrotestosterone | 3-oxo-5-beta-steroid 4-dehydrogenase |

| | | |
|---|---|---|
| DB07557 | 3,20-Pregnanedione | Nuclear receptor coactivator 1, Retinoic acid receptor RXR-alpha, Nuclear receptor subfamily 1 group I member 3, 3-oxo-5-beta-steroid 4-dehydrogenase |
| DB07717 | CYCLOPENTA[A]PHENANTHREN-3-ONE | Androgen receptor |
| DB08510 | 5-ALPHA-PREGNANE-3-BETA-OL-HEMISUCCINATE | Ig gamma-2 chain C region |
| DB08956 | Hydroxydione | |
| DB11371 | Alfaxalone | |
| DB11859 | Brexanolone | GABA(A) Receptor |
| DB12308 | Eltanolone | |
| DB12350 | Rostafuroxin | |
| DB12655 | Patidegib | G protein-coupled receptor Smoothened (Smo) inhibitor with antineoplastic activity |
| DB12972 | Sepranolone | |
| DB13012 | AQX-1125 | |
| DB13587 | Mesterolone | |
| DB13710 | Metenolone | |
| DB13951 | Stanolone acetate | Androgen receptor, Estradiol 17-beta-dehydrogenase 1, Estrogen receptor alpha, Mineralocorticoid receptor |
| DB14655 | Drostanolone propionate | |
| DB15490 | Zuranolone | GABA(A) Receptor |
| DB15830 | Ruscogenin | ruscogenin inhibited activation of neutrophil through cPLA2, PAK, Akt, MAPKs, cAMP, and PKA signaling pathways |
| DB16244 | MK-0773 | |
| DB16928 | Sarsagenin | |
| DB17016 | MK-4541 | |
| DB18200 | Golexanolone | |

**Table S2.** Neighboring molecules and reported mechanisms of action from ChEMBL dataset

| ChEMBL ID | Common name | Mode of action/Target Name |
|---|---|---|
| CHEMBL710 | finasteride | Inhibitor 5α-reductase |
| CHEMBL207538 | Allopregnanolone | GABA PAM |
| CHEMBL1200969 | Dutasteride | Inhibitor 5α-reductase |
| CHEMBL1568698 | GANAXOLONE | GABA PAM |

Table S4. Target gene abbreviations

| **Target gene** | **Abbreviations** |
|---|---|
| GABRA1-6 | GABA receptor subunit alpha 1-6 |
| AR | Androgen Receptor |
| ESR | Estrogen Receptor |
| G6PD | Glucose-6-phosphate dehydrogenase |
| GABRB1-3 | GABA receptor subunit beta 1-3 |
| GABRD | GABA receptor subunit delta |
| GABRE | GABA receptor subunit epsilon |
| GABRG1-3 | GABA receptor subunit gamma1-3 |
| GABRP | GABA receptor subunit pi |
| GABRQ | GABA receptor subunit theta |
| GRIN1 | glutamate ionotropic receptor NMDA type subunit 1 |
| GRIN2A/ GRIN2B/ GRIN2C/ GRIN2D/ GRIN3A/ GRIN3B | glutamate ionotropic receptor NMDA type subunit 2A/2B/2C/2D/3A |
| HSD17B1 | 17β-Hydroxysteroid dehydrogenase 1 |
| NR1I2/NR1I3 | nuclear receptor subfamily 1, group I, member 2/member 3 |
| PPARA | Peroxisome proliferator-activated receptor alpha |
| SIGMAR1 | sigma-1 receptor |

| | |
|---|---|
| SULT2A1, SULT2B1 | sulfotransferase family 2A member 1, sulfotransferase family 2B member 1 |
| GABRA | GABA receptor A |
| CXCR3 | C-X-C motif chemokine receptor 3 |
| VEGFR2 | vascular endothelial growth factor receptor 2 |
| Src, | non-receptor tyrosine kinase |
| FAK | focal adhesion kinase |
| CYP19A1 | cytochrome P450 family 19 subfamily A member 1 |
| ATP1A1 | Sodium/potassium-transporting ATPase subunit alpha-1 |
| CHRM2, CHRM3 | muscarinic acetylcholine receptor M2, M3 |
| CHRNA1, CHRNA2 | Neuronal acetylcholine receptor subunit alpha-1/2 |
| CYP17A1 | cytochrome P450 family 17 subfamily A member 1 |
| HSD3B1 | 3beta-hydroxysteroid dehydrogenase/delta(5)-delta(4)isomerase |
| OSC | oxidosqualene cyclase |
| GR | glucocorticoid receptor |
| HSD17B11 | Estradiol 17-beta-dehydrogenase 11 |
| IGHG2 | Ig gamma-2 chain C region |
